# Developmental dynamics of the neural crest-mesenchymal axis in creating the thymic microenvironment

**DOI:** 10.1101/2021.11.08.467624

**Authors:** Adam E. Handel, Stanley Cheuk, Fatima Dhalla, Stefano Maio, Tania Hübscher, Ioanna Rota, Mary E. Deadman, Olov Ekwall, Matthias Lütolf, Kenneth Weinberg, Georg Holländer

**Author notes:** These authors contributed equally (Adam E. Handel, Stanley Cheuk).

## Abstract

The thymic stroma is composed of epithelial and non-epithelial cells that collectively provide separate microenvironments controlling the homing of blood-born precursors to the tissue, and their subsequent differentiation to functionally mature and correctly selected T cells. While thymic epithelial cells are well characterized for their role in thymopoiesis, a comparably comprehensive analysis of the non-epithelial thymic stroma is lacking. Here we explore at single cell resolution the complex composition and dynamic changes that occur over time in the non-epithelial stromal compartment. We detail across different developmental stages in human and mouse thymus, and in an experimental model of Di George syndrome, the most common form of human thymic hypoplasia, the separate transcriptomes of mouse mesothelium, fibroblasts, neural crest cells, endothelial and vascular mural cells. The detected gene expression signatures identify novel stromal subtypes and relate their individual molecular profiles to separate differentiation trajectories and functions. Specifically, we demonstrate an abundance and unprecedented heterogeneity of diverse fibroblast subtypes that emerge at discrete developmental stages and vary in their expression of key regulatory signalling circuits and components of the extracellular matrix. Taken together, these findings highlight the dynamic complexity of the non-epithelial thymus stroma and link the cells’ specific gene expression profiles to separate instructive roles essential for normal thymus organogenesis and tissue maintenance.

**Teaser:** Single cell profiling of thymic stroma identifies a dynamic contribution from neural crest cells to the thymic mesenchyme.

## Introduction

Thymic T cell lineage commitment, development, maturation, and repertoire selection are instructed by a stromal scaffold that includes thymic epithelial cells (TEC) (*1*), endothelial cells (*2*) and mesenchymal cells (*3, 4*). The TEC compartment is both phenotypically and transcriptionally well characterized providing at single cell resolution a detailed account of the cells’ developmental dynamics and functions (*5, 6*). In addition, the thymus microenvironment is also composed of stromal cells of mesenchymal origin, including fibroblasts, endothelial cells and vascular mural cells. Derived primarily from either mesoderm or ectodermal neural crest cells, these thymic mesenchymal cells interact with TEC and thus create unique cellular niches that control thymopoiesis. This critical function of mesenchymal cells is accomplished via the production of extracellular matrix components, morphogens and key growth factors (*3, 4, 7*). Hence, the thymic mesenchyme is indispensable for the organ’s correct formation and function (*1, 7, 8*).

Fibroblasts constitute the largest component of the non-epithelial thymus stroma. (NETS) Though first described as distinct cell type over 150 years ago, the specific contributions of fibroblasts to organ formation, maintenance and function have only recently begun to be unraveled (*8*). Utilizing single-cell genomic technologies for the comparison of diverse tissues, fibroblasts were noted to display a significant heterogeneity with both cross-organ communalities and tissue-specific differences (*9*). Likewise, endothelial cells and vascular mural cells display organotypic features that have only recently been appreciated when resolving the cells’ distinct transcriptomes at single-cell resolution (*10*). In addition to their essential role in providing oxygen, nutrients, cells and other cargo to tissues, blood vessels also express in a context-specific fashion diverse transcriptomic profiles that include sets of growth factors inducing, specifying, patterning, and guiding organ formation and homeostasis (*11*). A third stromal component of non-epithelial origin are neural crest cells which enter the anlage as a migratory population as early as embryonic day 12 where they differentiate into distinct cell types, including vasculature-associated pericytes juxtaposed between endothelia and the other components of the stromal scaffold (*12*).

A detailed phenotypic, transcriptomic, and functional genomic description of the diverse population of non-epithelial thymus stromal cells is to date still wanting. We have therefore employed flow cytometry and single cell multiomics technologies to detail the complexity and developmental dynamics of thymic mesenchymal cells in both mouse and human tissue. Our results highlight a previously unappreciated heterogeneity among cells belonging to the NETS under physiological conditions and identify distinct yet selective defects of these cells in a genetic mouse model of the 22q11 deletion syndrome, the most common human condition associated with congenital thymus hypoplasia.

## Results

### Single cell sequencing reveals high levels of complexity within the thymic mesenchyme

We first sought to delineate both the frequency and diversity of NETS cells (phenotypically defined as Ter119^-^CD45^-^EpCAM^-^) in the thymus of 4-week-old mice. These cells accounted for approximately half of the total thymus stroma cellularity and distinct subpopulations were identified using the differential expression of glutamyl aminopeptidase Ly51, glycoprotein podoplanin (gp38) and dipeptidyl peptidase-4 (DPP4, CD26) (Fig. 1a and Fig. S1a) (*3, 13*). The Ly51^hi^gp38^-^ phenotype identified neural crest-derived pericytes which surround blood vessels adjacent to endothelial cells (*4, 12*). The gp38 positive stromal cells expressed a reduced level of Ly51 and could be further differentiated into separate subpopulations based on their CD26 expression: gp38^+^CD26^+^ cells localized to the thymus capsule whereas gp38^+^CD26^-^ were enriched in the medulla (Fig. 1b) (*13*).

**Fig. 1.**
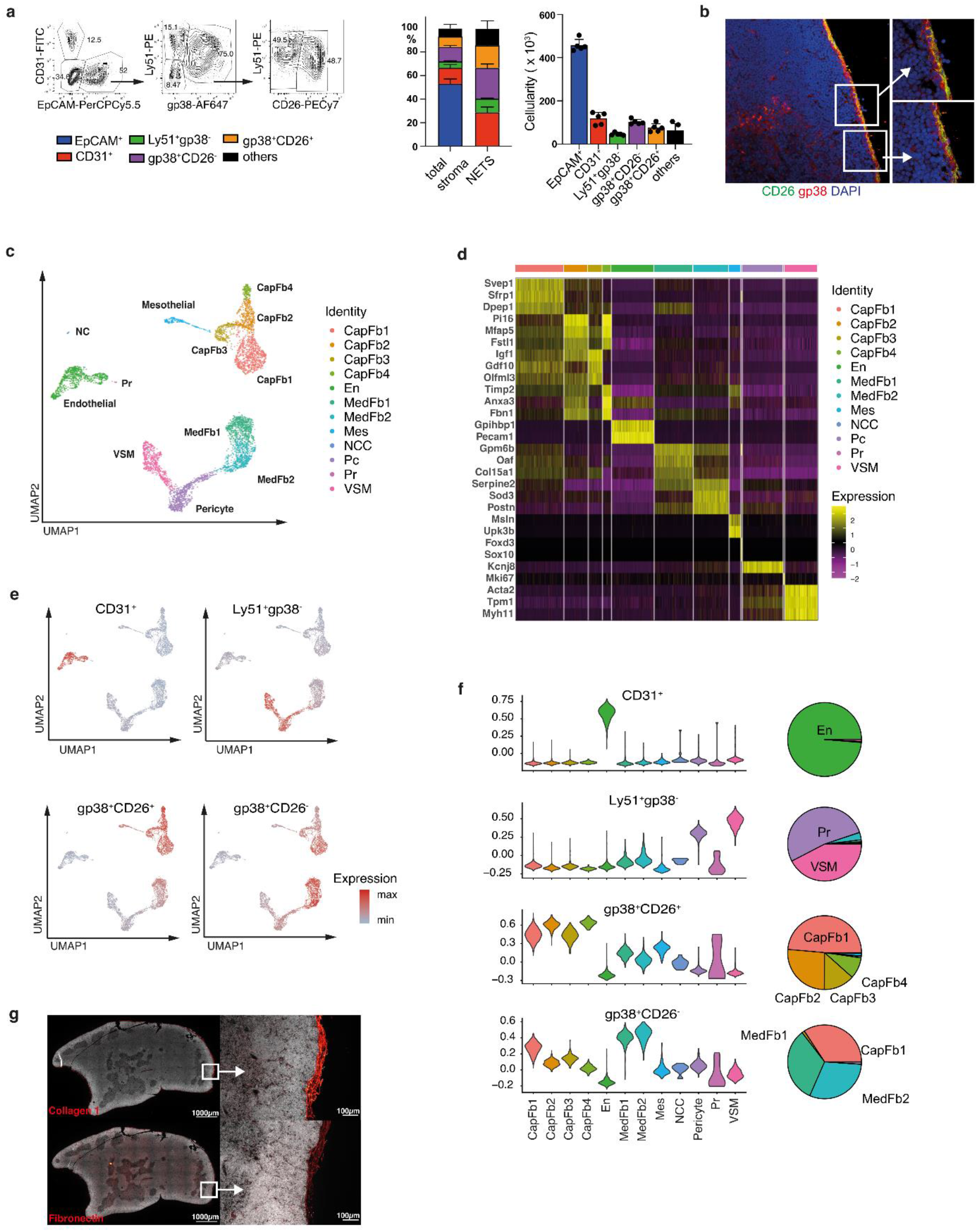
Heterogeneity of the thymic stroma at 4 weeks of age. (**a**) Representative FACS plot of live Ter119^-^ CD45^-^ EpCAM^-^ thymic stromal cells at 4 weeks old (left), the relative frequency (middle) and cellularity (right) of CD31^+^, Ly51+gp38^-^, CD26-gp38^+^ and CD26+gp38^+^ cells among total stroma or non-epithelial stroma. (b) Immunofluorescence staining of thymic mesenchymal cells (Red: gp38, Green: CD26, Blue: DAPI). (**c**) A UMAP plot of Ter119-CD45-EpCAM-cells from 4-week-old mice. (**d**) A heatmap of top 5 differentially expressed genes between each cluster. (**e**) An overlay on UMAP plot and (**f**) violin plots of the expression of genes specific to each FACS-isolated subpopulation from bulk RNA-seq and pie charts showing the proportion of cell types which express gene signatures characteristic of specific FACS-isolated populations as inferred from bulk RNASeq. CapFb = capsular fibroblast; En = endothelium; MedFb = medullary fibroblast; Mes = mesothelium; NCC = neural crest; Pc = pericyte; Pr = proliferating cell; VSM = vascular smooth muscle. (**g**) Immunofluorescence microscopy showing the distribution of type-I and fibronectin in a 5-week-old mouse thymus. FACS data shown in panel a were representative of one experiment (n=5) out of two independent experiments (total n=7), and mean value and SD are shown in the corresponding bar graphs (n=5).

To delineate the heterogeneity of the thymic mesenchyme in an unbiased fashion and independent of a limited number of phenotypic markers, we next generated transcriptomic libraries from 5,878 single Ter119^-^CD45^-^EpCAM^-^ cells isolated from 4-week old thymi (Extended Data Fig. 1a). We identified 12 distinct cell subtypes based on their separate gene expression profiles (Fig. 1c, d, Fig. S1b). (For clarity, we refer to transcriptionally defined stromal cell clusters as subtypes whereas the terms populations and subpopulations specify cells that have been defined by cytometry). A Uniform Manifold Approximation and Projection (UMAP) analysis of non-TEC stroma identified four separate cell clusters of different sizes. The two largest comprised several individual cell subtypes that were transcriptomically defined as either capsular or medullary fibroblasts (Fig. 1c and Fig. S1c i and ii) (*13*). The capsular cluster closely resembling the gene expression profile of capsular fibroblasts consisted of four subsets that separated from mesothelial cells defined – *inter alia* - by their expression of *Msln* and *Upk3b*, encoding the glycosylphosphatidylinositol-anchored cell-surface adhesion protein mesothelin and the membrane integral protein Uroplakin, respectively (*14*). Within the capsular fibroblast clusters, subtype 1 (designated CapFb1) was characterized by the high expression of *Svep1, Sfrp1* and *Dpep1*, which encode a multidomain cellular adhesion molecule (*15*), the secreted frizzled-related protein 1 modulating stromal to epithelial signaling via Wnt inhibition (*16*), and a membrane-bound dipeptidase involved in the metabolism of glutathione and other similar proteins (*17*). The capsular subtypes 2 (CapFb2) and 4 (CapFb4) were characterized by the expression of *Pi16, Mfap5* and *Fstl1*, which encode a peptidase inhibitor of largely unknown function, the microfibrillar-associated protein 5 related to extracellular matrix remodeling and inflammation (*18*), and the secreted extracellular glycoprotein follistatin-like 1. CapFb4 also highly expressed *Timp2* encoding the tissue inhibitor of metalloproteinase 2 relevant for tissue remodeling (*19*), *Anxa3* translating into the membrane-associated Annexin 3 protein activating the epithelial-to mesenchymal transition (EMT) program and Wnt signaling pathway (*20*), and *Fbn1* encoding Fibrillin 1, a major component of extracellular microfibrils. The CapFb3 subtype typically expressed *Igf1* encoding Insulin-growth factor 1 regulating tissue homeostasis via cell proliferation, differentiation, maturation, and survival (*21*), *Gdf10* translating into the transforming growth factor-β (TGF-β) superfamily member growth differentiation factor-10 (GDF10) (*22*), and *Olfml3* encoding the secreted glycoprotein olfactomedin-like 3 which has matrix-related functions central to embryonic development (*23*).

The second UMAP cluster incorporated two distinct medullary fibroblast subtypes, pericytes and vascular smooth muscle cells. The medullary fibroblast subtypes 1 (MedFb1) and 2 (MedFb2) displayed similar gene expression profiles although transcripts for the out-at-first protein (encoded by *Oaf*) and the alpha 1 chain of collagen XV (*Col15a1*) were detected at higher levels in MedFb1 while transcripts for extracellular superoxide dismutase 3 (*Sod3*) were particularly evident in MedFb2. Both subtypes also comprised transcripts for IL-33 and Cxcl16, which are important for dendritic cell activation and NKT cell migration, respectively (*24, 25*). Transcripts related to antigen processing and presentation were enriched in MedFb2 (Fig. S1b). However, contrary to a recent observation (*13*), tissue restricted antigens were not generally more frequent in medullary fibroblasts when compared to other thymic NETS (Fig. S1e).

Pericytes were identified by their characteristic expression of *Kcnj8* encoding member 8 of the J subfamily of the potassium inwardly rectifying channels, which form part of the ATP/ADP-binding potassium channel of these cells (*26*). Vascular smooth muscle cells (VSM) were characterized by the expression of contractile elements, including *Acta2, Tpm1* and *Myh11*, whereas endothelial cells displayed a high number of transcripts for *Gpihbp1* and *Pecam1* which encode the glycosylphosphatidylinositol anchored high density lipoprotein binding protein 1 and the intercellular junction protein platelet and endothelial cell adhesion molecule (PCAM, aka CD31), respectively. Neural crest-derived cells (NCC) were characterized by their expression of *Foxd3* and *Sox10* which are critical for the cells’ specification and development (*27, 28*). Finally, actively proliferating cells were identified by the expression of different cell cycle-related genes including *Mki67* encoding the nuclear protein Ki67.

We determined the gene expression profiles of the 4 FACS-defined non-TEC thymic stromal subpopulations (Fig. S1a) and deconvoluted the individual transcriptomes by projection onto the single cell UMAP data (Fig. 1e, Table 1). Stroma cells expressing CD31 identified the cluster defined as endothelial cells and the Ly51^+^Gp38^-^ subpopulation represented the pericyte and vascular smooth muscle cell clusters (Fig. 1e, f). The gp38^+^CD26^+^ subpopulation included all of the 4 capsular fibroblasts subtypes, whereas the gp38^+^CD26^-^ subpopulation was mainly enriched for the medullary fibroblast subtypes but also included CapFb1 cells (Fig. 1e, f). We showed that this fibroblast heterogeneity patterned the thymic extracellular matrix by staining for two key extracellular matrix molecules (type I collagens and fibronectin), which were most highly expressed in capsular fibroblasts (Fig. 1g).

**Table 1.**
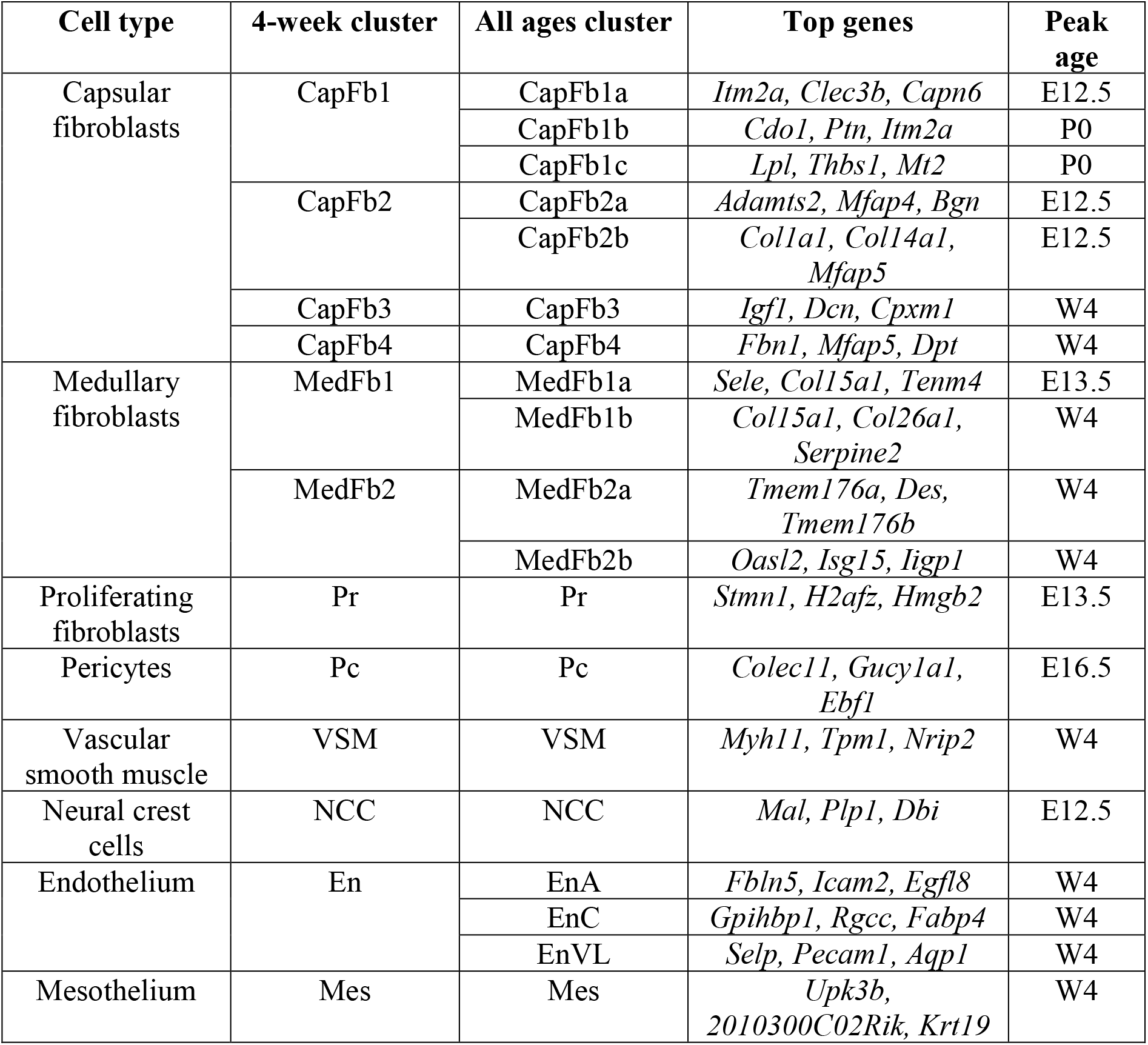
Thymic stromal cell types identified by single cell RNA-seq. The three top genes (by area under the curve) and peak age are shown for each cluster. All clusters show significant differences in proportional makeup of the non-TEC thymic stromal across different ages (Fisher’s test with 10,000 permutations: all p < 0.0001).

Hence, the single cell RNA-seq based identification of thymic stromal cells unmasked a previously unrecognized heterogeneity of individual subsets among gp38^+^ NETS which could not be identified by conventional flowcytometry-based phenotyping.

### Thymic organogenesis is characterised by dynamic mesenchymal changes

The thymus undergoes significant micro-architectural changes during organogenesis, including the compartmentalisation into distinct cortical and medullary domains and the formation of a complex vascular network (*29*). We therefore investigated how these morphological changes paralleled compositional alterations of the mesenchymal stroma (Fig. 2a-b). At embryonic day (E)12.5, NETS accounted for more than 90% of all CD45^-^ thymic cells with Ly51^+^gp38^-^ cells being by far the most dominant subpopulation. The frequency of TEC gradually increased parallel to thymus growth and reached a relative maximum at E16.5 when epithelia represented 60% of the thymic stroma. Earlier during thymus organogenesis, the non-TEC stroma lacked the heterogeneity observed at E16.5 and thereafter. For example, gp38^+^CD26^-^ and gp38^+^CD26^-^ fibroblasts dominated the stromal compartment at both E12.5 and E13.5, endothelial cells were only discovered at E13.5 and Ly51^+^gp38^-^ pericytes were not detected prior to E16.5 (Fig. 2a, Fig. S2a). The subpopulation of gp38^+^CD26^+^ capsular fibroblasts was identified as early as E13.5 and increased in frequency thereafter (Fig. 2a).

**Fig. 2.**
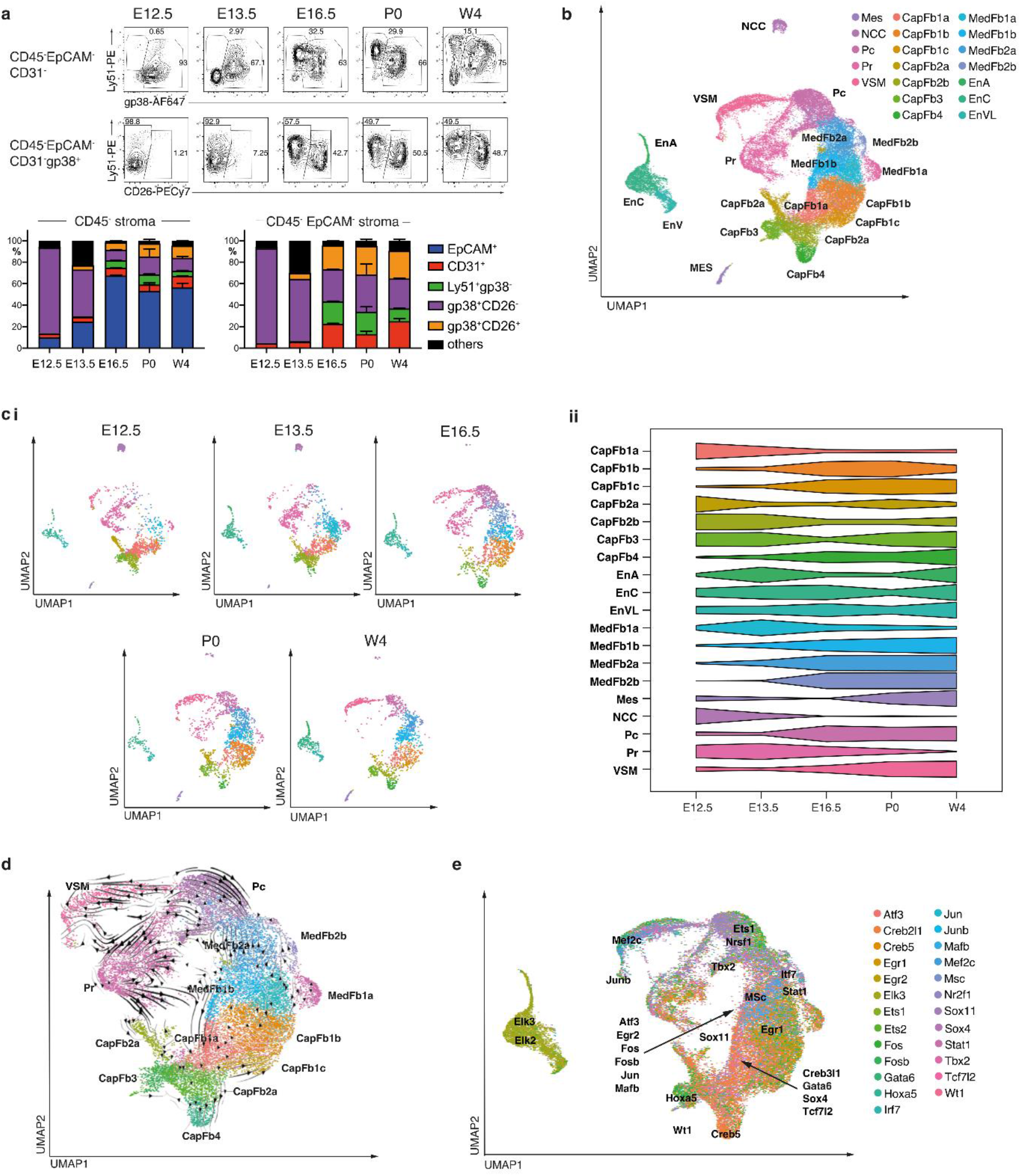
Characteristics of the thymic stroma over developmental time. (**a**) (upper) representative FACS plot of the non-epithelial thymic stroma from E12.5 to week 4. (lower) and the relative frequency of stromal subpopulations among the total stroma (left) or the non-TEC stroma (right). (**b**) A combined UMAP plot of Ter119-CD45-EpCAM-cells from mice at E12.5, E13.5, E16.5, P0 and 4 weeks of age. (**c**) (i) Individual UMAP plots for each developmental timepoint, with cell number down-sampled to the smallest sample size (n = 1,997). (**ii**) Scaled proportional representation of each age in each cluster. (**d**) A UMAP plot with each cell coloured by the maximum transcription factor gene-regulatory network expression. Gene-regulatory network centroids are labelled. (**e**) An RNA velocity plot with velocity streamlines projected onto UMAP plot. CapFb = capsular fibroblast; EnA = arterial endothelium; ENCC = capillary endothelium; EnVL = venous/lymphatic endothelium; MedFb = medullary fibroblast; Mes = mesothelium; NCC = neural crest; Pc = pericyte; Pr = proliferating cell; VSM = vascular smooth muscle. E12.5 and E13.5 were sorted and analysed from thymi pooled from two litters consist of at least 3 embryos per litters. For E16.5, P0 and week 4, data shown consist of cells sorted from two thymi per timepoint. Mean value and SD were shown in the bar charts (a).

We next used single cell RNA-seq to detail changes in the heterogeneity of individual NETS subtypes and to determine the cells’ developmental trajectories. We generated libraries on a total of 36,208 single stromal cells isolated from embryonic (E12.5, 13.5, 16.5), new-born and young adult thymus tissue, which collectively reiterated the clusters observed in the thymus of 4-week-old mice and provided sufficient resolution to identify additional heterogeneity (Fig. 2b-d, Table 1). Complex dynamic changes in the frequency of individual subtypes occurred over time between the early developmental stages and the completion of a mature thymus microenvironment. For example, CapFb1a and CapFb2b appeared early but their frequencies gradually decreased during organogenesis whereas all of the medullary fibroblasts (with the notable exception of MedFb1a) increased parallel to the emergence of mTEC (*30*). NCC were largely absent after E16.5 but other NETS subtypes remained either mainly unchanged or displayed a bi-model variation in frequency between E12.5 and 4 weeks of age (Fig. 2c i and ii). This finding is in agreement with lineage tracing studies demonstrating the cells’ developmental potential to differentiate into VSM and pericytes.(*12, 31*) Thus, single cell RNA-seq revealed complex and dynamic changes in the relative number of individual NETS subtypes that would be captured incompletely by classical cell surface phenotyping, such as CD26 (Fig. S2b).

We leveraged the splicing information obtained from single cell transcriptomes to determine the developmental trajectories of individual NETS subtypes. This analysis identified the CapFb1a and CapFb2b subtypes as the principal precursors for other capsular fibroblasts (Fig. 2d) and suggested MedFb1a to serve as a precursor for other fibroblast subtypes in the emerging medulla (*30*). This analysis also recognized CapFb3 fibroblasts as intermediates between mesothelial cells and other fibroblast subtypes, a finding consistent with the concept that fibroblasts can be derived from mesothelial cells (*32*). However, CapFb3 were distinct from mesothelia as they lacked the expression of *Msln* and *Upk3b* (Fig. 1d) (*14*).

The single cell transcriptome data was also used to infer gene regulatory network activities of individual NETS subtypes (Fig. 2e). A transcription factor motif analysis of these gene regulatory networks was executed to identify potential cell type-specific transcription factors (*33*). In keeping with their proposed differentiation from mesothelial cells, CapFb3 fibroblasts expressed gene regulatory networks controlled by the transcription factors *Hoxa5* and *Wt1*, which typically are active in mesothelial cells (Fig. 2e, Fig. S2c) (*14*). CapFb4 were highly enriched for a *Creb5*-controlled gene regulatory network that has previously been identify to modulates the differentiation of fibroblasts to myofibroblasts (*34*) and to control age-related thymic fibrosis (*35*) (Fig. S2d). MedFb2b expressed *Irf7* encoding the Interferon-regulatory factor 7 (IRF7), a master regulator of type I IFN secretion that interacts with Smad3 to regulate TGF-β signalling for collagen production (Fig. 2e, Fig. S2e) (*36*).

We assessed the expression of canonical Wnt signalling transcripts and growth factors known to be important in thymic stromal interactions with thymocytes (Fig. S3) (*37*). Several Wnt ligands displayed distinct expression patterns among cells of the NETS. For example, *Wnt4* transcripts were detected in mesothelium, *Wnt5a* in CapFb2b, CapFb3 and CapFb4, *Wnt6* in NCC, and *Wnt10b* in CapFb4. Wnt modulators were also highly expressed in particular non-epithelial stroma cells, including *Rspo1* in mesothelium and *Sfrp5* in NCC. Several cell subtypes of the NETS acted as prominent sources of key growth factors, including *Bmp4* transcribed in CapFb1c, *Bmp7* in CapFb4 and mesothelium, *Fgf10* in CapFb1a, CapFb1b and CapFb1c, and *Tgfb1* in endothelial cells. In keeping with their role in regulating the extracellular matrix, fibroblast subtypes showed high expression of key extracellular matrix transcripts, with collagens (*e*.*g. Col1a2, Col3a1* and *Col14a1*) primarily expressed in capsular fibroblasts, and laminins (*Lama2* and *Lama4*) in a mixture of capsular (CapFb1a, CapFb1b and CapFb1c) and medullary fibroblasts (MedFb1b and MedFb2b). This distinctive expression of growth and differentiation factors, and components of the extracellular matrix demonstrated that heterogeneity within the NETS compartment determined modularity in the expression of key molecules, thus implicating different developmental and functional niches.

### Ligand-receptor pairing analysis identifies interactions between neural crest-derived mesenchyme and endothelial cells

Given that NCCs are known to differentiate into perivascular cell types, we aimed to uncover the ligand-receptor signaling and the subsequent transcriptomic networks that control the differentiation of NCCs into pericytes and VSM and pericytes (*12, 31*). To this end, NicheNet identified intercellular ligand-receptor interactions associated with cell type-specific transitions across early (E12.5 and E13.5) to later stages (E16.5) in embryonic thymus formation (*38*). This analysis demonstrated that ligands expressed by endothelial cells, including the adhesive and multimeric glycoprotein von Willebrand factor (vWF) and Transforming Growth Factor Beta 1 (TGFB1), influenced gene expression in thymic NCC (Fig. 3a, b). Conversely, heterotypic interactions between junctional adhesion molecule-3 (*Jam3*) produced by NCC and its receptor *Jam2* on endothelial cells identified a candidate ligand-receptor pair that orchestrated the changes in the gene expression profile of embryonic endothelial cells (Fig. 3c). Together, these inferred ligand-receptor interactions suggested that reciprocal cellular relationships between vascular structures and NCC shape the perivascular thymic stroma during embryogenesis.

**Fig. 3.**
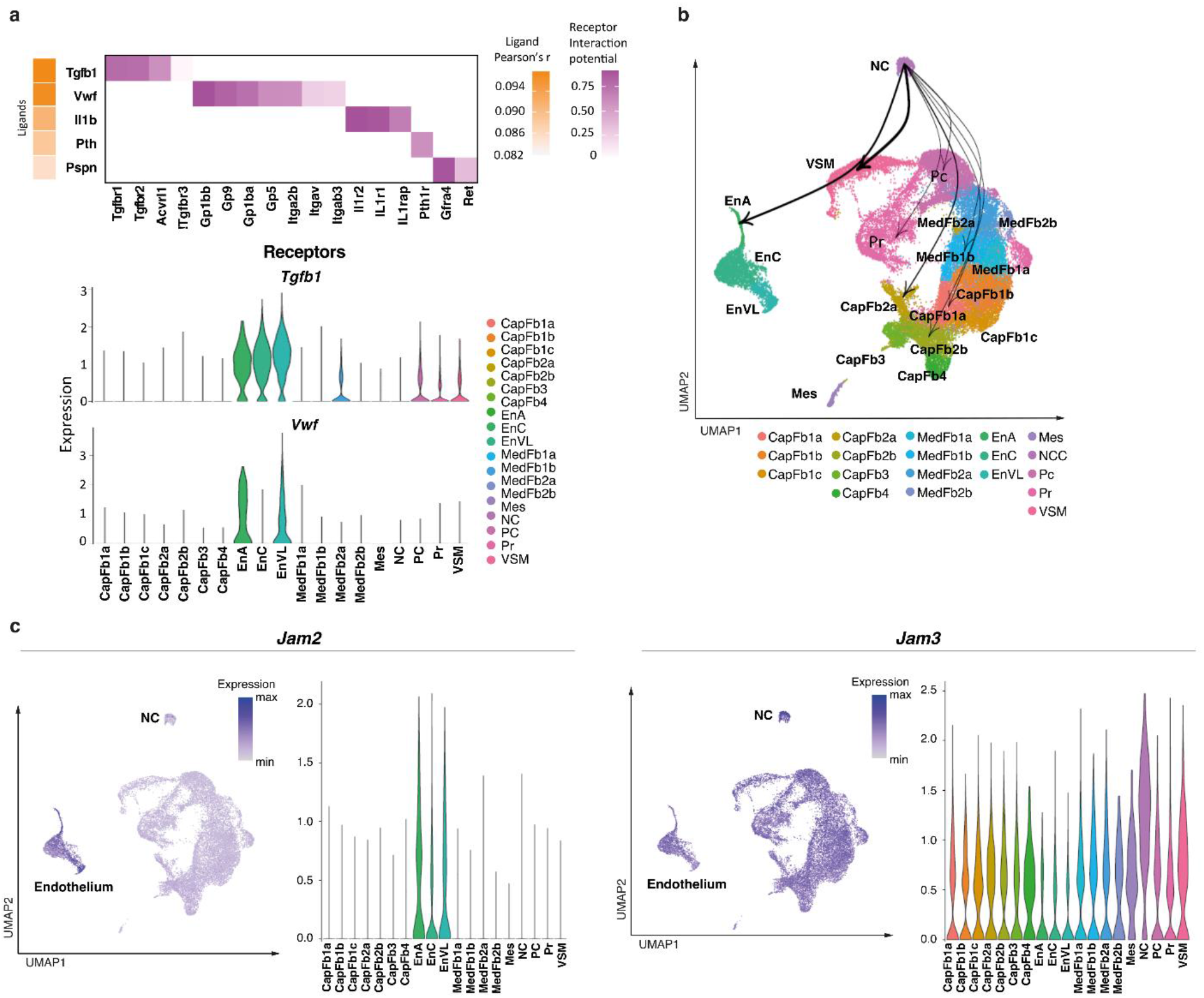
Ligand-receptor interaction of neural crest derived cells and other thymic cell types. **(a) (Upper)** Heatmap of top ligand-receptor interactions of cell types signalling to NCC showing the Pearson’s r for ligand activity in promoting ageing transition between E12.5/13.5 and E16.5. **(Lower)** Violin plots of the cell type-specific expression of the top two ligands, Tgfb1 and Vwf. **(b)** A UMAP plot showing weighted connections for the top 25% ligand-receptor-target networks for connections from neural crest cells. The width of the line is proportional to the strength of ligand-receptor interactions. **(c)** UMAP and violin plots of *Jam2* and *Jam3* expression.

### Reduced cellularity and complexity of mesenchymal stroma are features of an experimental 22q.11.2 Deletion Syndrome model

A spontaneously occurring heterozygous deletions of 1.5Mb to 3Mb size following recombination between four blocks of low copy repeats within 22q11.2 account for the loss of up to 106 genes (*39*). This mutation constitutes the most common molecular etiology of the 22q11.2 deletion syndrome (DS; previously referred to as DiGeorge Syndrome (DGS)) which manifests clinically as a range of features that include either athymia resulting in T cell deficiency or thymus hypoplasia compromising immunological fitness (*39*). Regions of mouse chromosome 16 are syntenic to the human 22q11.2 (*40*) and include *Tbx1*, encoding a T-box transcription factor and *Crkl*, encoding an adapter protein implicated in fibroblast growth factor and focal cell adhesion signaling (*39*). Compound haploinsufficiency of *Tbx1* and *Crkl* in mice results in typical hallmarks of 22q11.2DS, including thymic hypoplasia (*41*).

The abnormal migration of cephalic NCC has been identified as a possible cause for the pharyngeal patterning defects observed in 22q11.2DS which is recapitulated in mice compound heterozygous for *Tbx1* and *Crkl* (designated *Tbx1*^+/-^*Crkl*^+/-^). Gene products of these two loci have been alleged to interact in a dosage-sensitive fashion (*41*). *Tbx1* expression in NETS was exclusively confined to E12.5 and detected in a subset of cortical fibroblast subtypes, especially CapFb4, and proliferating cells. Yet, *Crkl* transcripts were mainly detected in CapFb1 and CapFb4 subtypes early during thymus development but could also be identified in a small fraction of these and other NETS later in development (Fig. S4a-c).

The thymi of mice compound heterozygous for a loss of *Tbx1* and *Crkl* were hypoplastic and revealed already at E13.5 significantly fewer haematopoietic, epithelial and mesenchymal cells than their wild type controls, which was not the case for either single *Tbx1* or *Crkl* mutants (Fig. 4a-c and Fig. S4d). At birth, hematopoietic cells and all phenotypically identified major NETS subpopulations were reduced in mutant mice and remained diminished in 4-week-old mice with the notable exception of gp38^+^CD26^+^ capsular fibroblasts. In contrast, the cellularity of TEC and endothelial cells varied over time and were not uniformly reduced in mutant mice at these times (Fig. 4b, c; Fig. S4e). Thus, several NETS subpopulations were consistently reduced in mice heterozygous for *Tbx1* and *Crkl*.

**Fig. 4.**
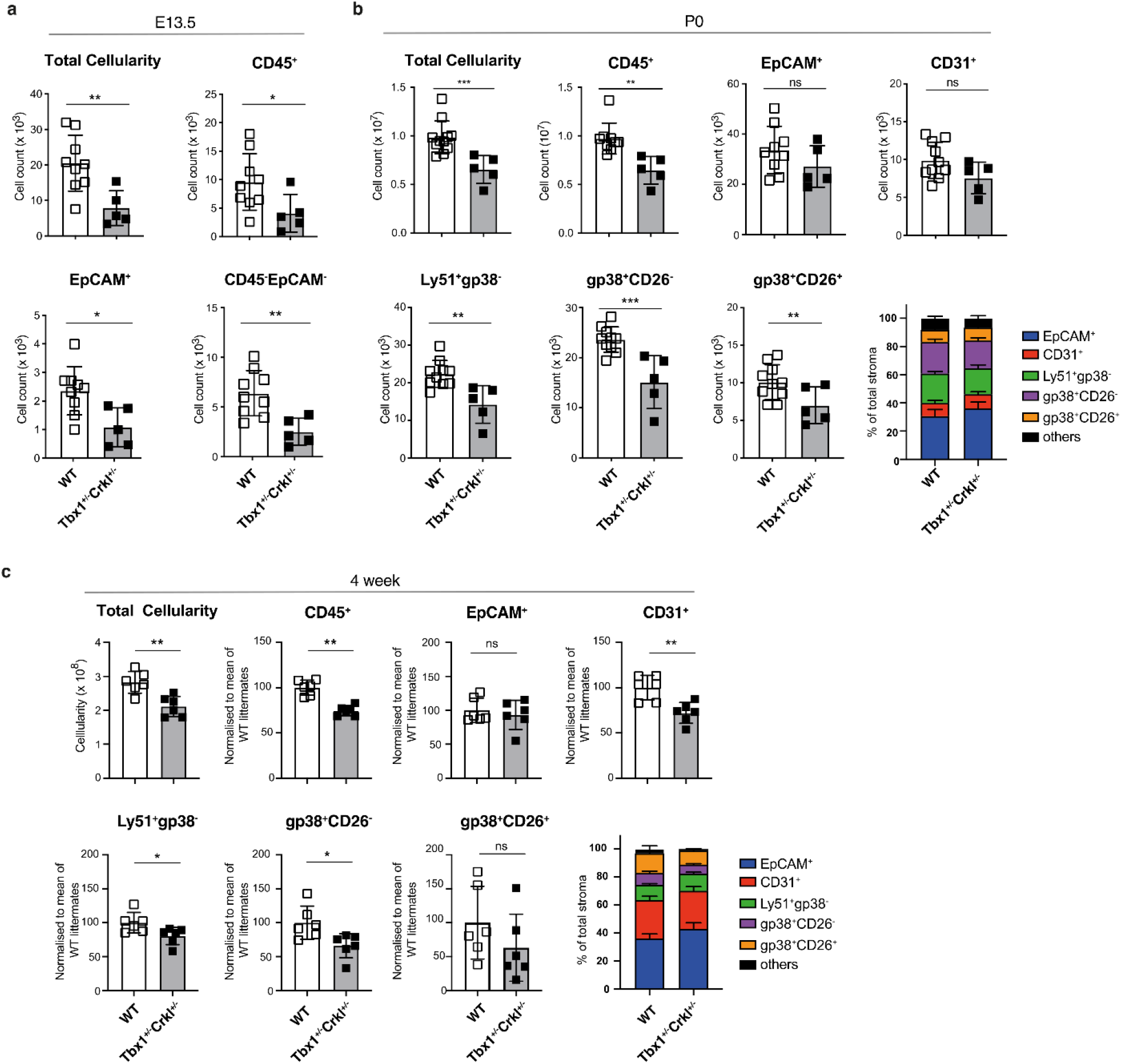
Reduction of non-epithelial thymic stroma in Tbx1+/-Crkl+/-thymi. **(a)** Total cellularity, Number of CD45^+^, EpCAM^+^ and total mesenchymal cells at E13.5 wildtype (n=9) and Tbx1^+/-^Crkl^+/-^ (n=5) thymi. **(b)** Total thymic cellularity, absolute number of CD45+, EpCAM+, CD31+, Ly51^+^gp38^-^, CD26^-^gp38^+^, CD26^+^gp38^+^ cells and the frequency of stromal subpopulations from wildtype (P0: n=9, 4-week: n=6) and Tbx1^+/-^Crkl^+/-^ (P0: n=5, 4-week: n=6) neonates **(b)** and 4-week old mice **(c)**. Data was normalised to the mean of wildtype litter mates from two independent experiments (n=3 from each experiment). Mean values were shown in the bar charts. Unpaired t-test, p<0.05(*), 0.01(**), 0.001(***).

We next compared the transcriptome of individual epithelial and NETS cells isolated from new-born *Tbx1*^*+/-*^*Crkl*^*+/-*^ mice (Fig. 5a and Fig. S5a; epithelial and non-epithelial stromal cells were annotated as previously published (*5*) and shown in Figure 2, respectively). *Tbx1* and *Crkl* compound heterozygosity substantially changed the composition of the thymus stroma resulting in a reduction of 8 of the 18 individual subtypes in *Tbx1*^*+/-*^*Crkl*^*+/-*^ mice. Specifically and in contrast to the results obtained by flow cytometry, the relative cellularity of several capsular and medullary fibroblast subtypes together with that of pericytes and VSM was lessened (Fig. 5b). In addition, the frequencies of mature cortical (mcTEC) and intertypical TEC (itTEC) were reduced in mutant mice whereas that of perinatal cortical TEC (pcTEC), post-Aire mTEC (pamTEC) and neural TEC (nTEC) were enriched (Fig. 5c).

**Fig. 5.**
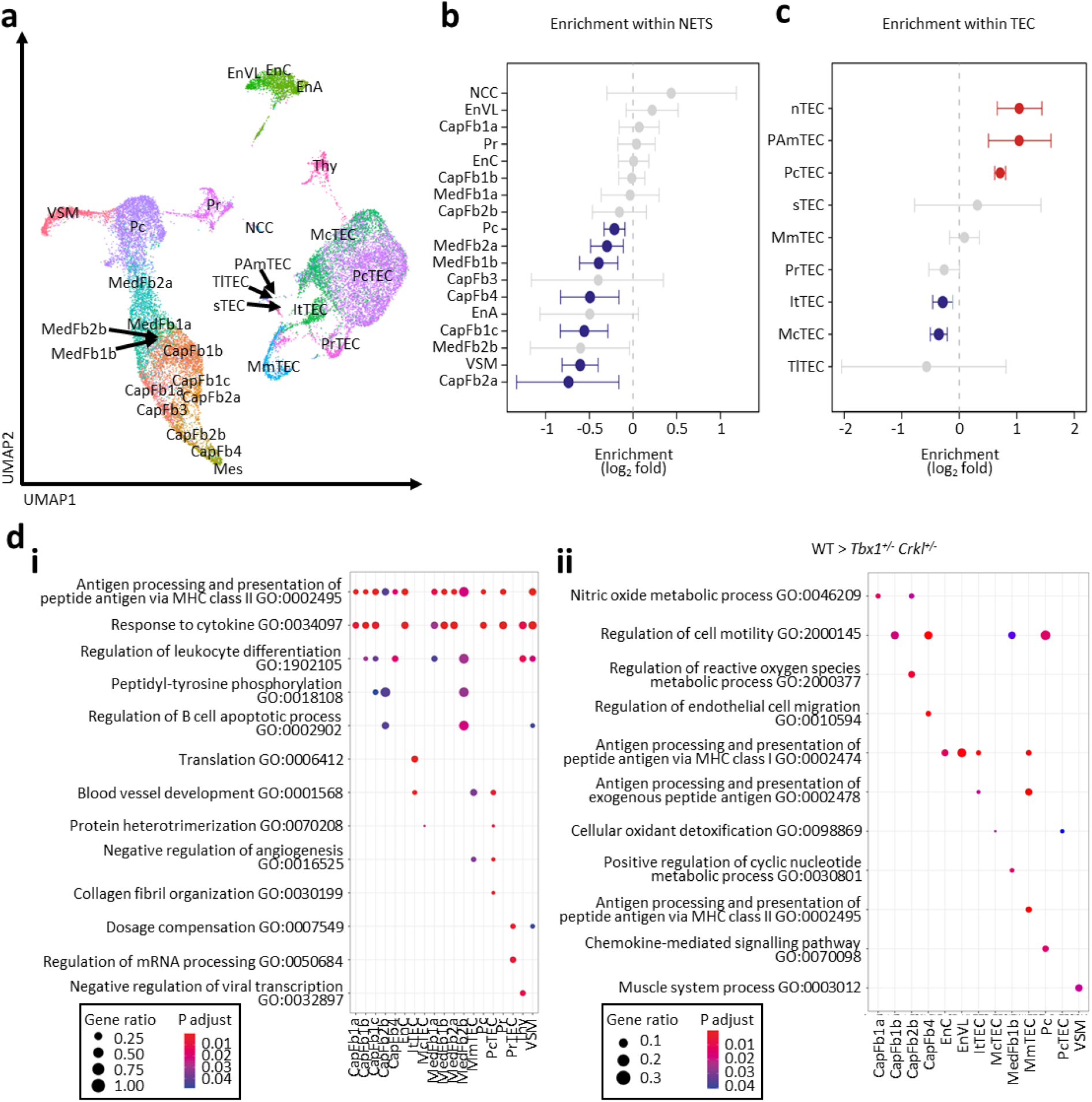
Thymic stromal cells from Tbx1^+/-^Crkl^+/-^and wild-type mice differ transcriptomically within cellular populations. A total of 10,300 cells from *Tbx1*^*+/-*^*Crkl*^*+/-*^ and 11,151 cells from their wildtype littermates were analysed. (**a**) A UMAP plot showing thymic stromal cell types. Scatter plot of genotype-specific enrichment of cell type frequency in Tbx1^+/-^Crkl^+/-^ and wild-type within NETS (**b**) and TEC (**c**). Error bars show 95% confidence intervals. (**d**) Gene ontology analysis of genes more highly expressed within each cell type in (**i**) Tbx1^+/-^Crkl^+/-^ and (**ii**) wild-type thymi. (e) Scatter plot of the difference in ageing score of stromal cell subsets between Tbx1^+/-^Crkl^+/-^ and wild-type. CapFb = capsular fibroblast; EnA = arterial endothelium; ENCC = capillary endothelium; EnVL = venous/lymphatic endothelium; itTEC = intertypical TEC; McTEC = mature cortical TEC; MedFb = medullary fibroblast; Mes = mesothelium; mmTEC = mature medullary TEC; NCC = neural crest; pAmTEC = post-AIRE medullary TEC; Pc = pericyte; pcTEC = perinatal cortical TEC; pr = proliferating cell; prTEC = proliferating TEC; sTEC = structural TEC; Thy = thymocyte; TlTEC = tuft-like TEC; VSM = vascular smooth muscle. Symbol in red showed cell subtypes significantly enriched in Tbx1^+/-^Crkl^+/-^ and symbol in blue showed population showed cell subtypes enriched in wildtype **(b**,**c)**. Enrichments were calculated using Fisher’s exact test with 95% confidence intervals and significance was adjusted for multiple hypothesis testing using Benjamini-Hochberg correction (b,c)

To assess the functional consequences of a compound heterozygous loss of *Crkl* and *Tbx1*, we performed an enrichment analysis for differentially expressed gene sets (Fig. 5d, Fig S5b). This analysis revealed that transcripts for gene products relevant for cell migration were reduced in several capsular and medullary fibroblast as well as in pericytes. In VSM, fewer transcripts for contractile elements (e.g. *Myl6* and *Tpm2*; Fig 5d) were observed.

### Compound *Tbx1* and *Crkl* heterozygosity is associated with accelerated aging of thymic mesenchyme

22q11DS patients display an accelerated thymic senescence (*42*), a process implicated to be caused by an age-dependent chronic systemic inflammatory condition known as inflammaging (*43*). To appraise the effects of inflammaging on NETS, we applied to our data an ageing score computed from age-driven transcriptomic changes common across many tissues (*44*). In contrast to the heterogeneous effect of ageing on TEC subsets (*5*), the ageing module score of non-TEC stroma progressively increased from early embryonic stages to young adulthood (Fig. 6a). These changes demonstrated a switch from an abundant expression of transcripts belonging to biosynthetic pathways to gene products associated with angiogenesis and immunological crosstalk (Fig. S6a).

**Fig. 6.**
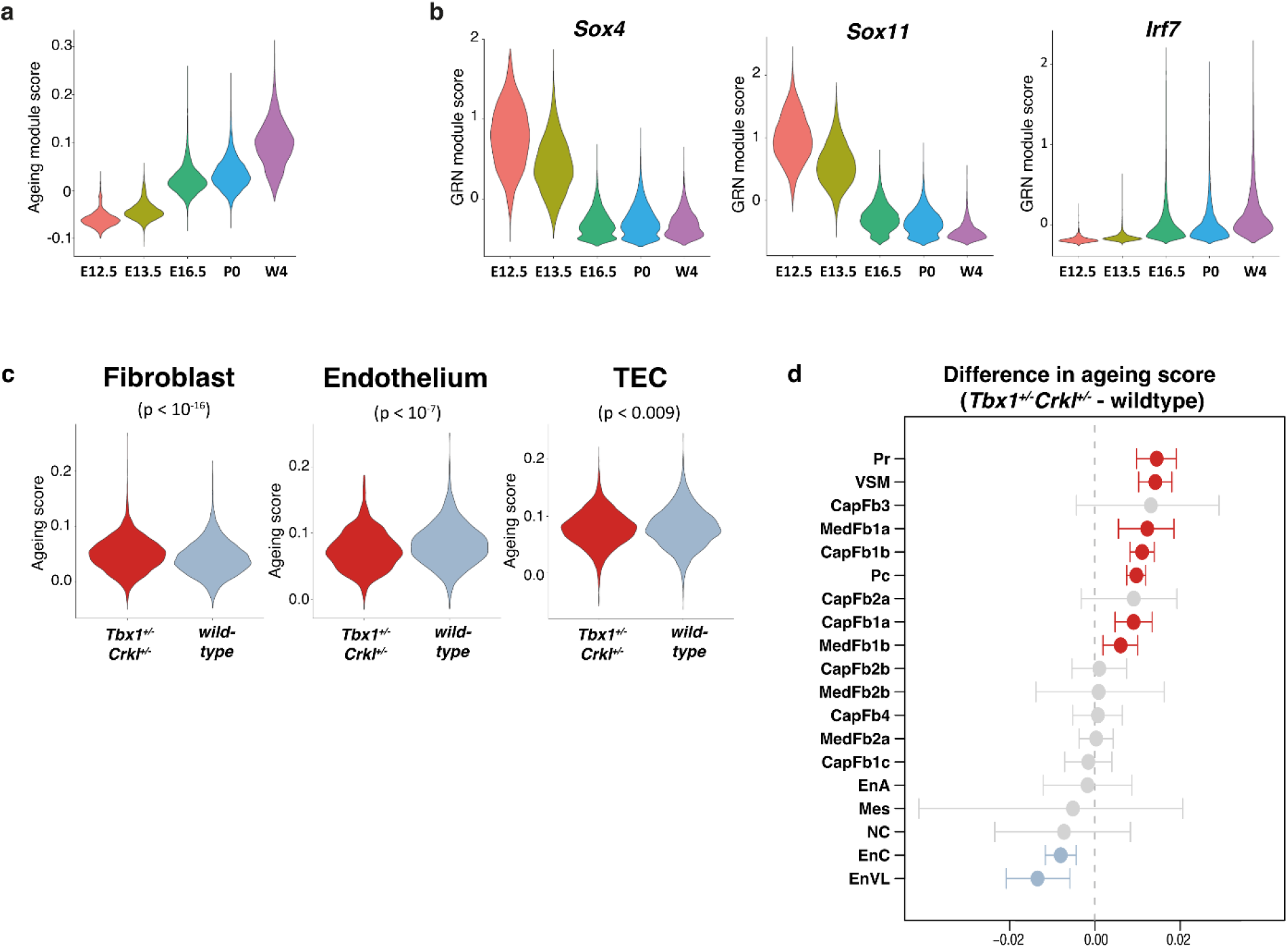
Age-specific transcriptomic programmes differ over development in thymic stroma. (**a**) A violin plot showing the overall expression of genes associated with ageing in a tissue-independent manner within the NES across different ages (*44*). (**b**) Violin plots showing the expression of Sox4, Sox11 and Irf7 gene-regulatory networks. **(c)** Violin plot showing the ageing score of fibroblasts, endothelial cells and TECs from Tbx1^+/-^Crkl^+/-^ and wild-type thymi at P0. Symbols in red showed cell subtypes significantly increased ageing score in Tbx1^+/-^Crkl^+/-^ as compared to wildtype and symbols in blue showed population showed significantly reduced ageing score **(d)**. Differences in gene module scores were estimated using Wilcoxon tests and were adjusted for multiple hypothesis testing using Benjamini-Hochberg correction (c,d).

Altered transcription factor network activity has been linked to the process of senescence (*44*). Using the transcriptomes from NETS isolated from mice at different ages, a decrease in transcripts related to gene networks controlled by Sox 4 and its close relative Sox11 were observed. In parallel, gene networks controlled by IRF7 were gradually activated over time and beyond what would be expected from their enrichment in MedFb2b subsets (Fig. 6b and Fig. S2e). In contrast, *Fos* and *Fosb* controlled gene networks peaked at birth but were subsequently weakened (Fig. S6b), a pattern previously noted in other tissues (*45*).

The ageing module score analysis was extended to include thymic stromal cells isolated from *Tbx1*^*+/-*^*Crkl*^*+/-*^ mice at postnatal day 0. Accelerated aging (as discernible by an increased score) was noted in mutant mice for the population of mesenchymal but not endothelial and epithelial cells (Figure 6c). Within the mesenchymal compartment, accelerated aging was not uniform as an increased score was observed in only 7 of the 19 distinct stromal subtypes including pericytes, VSM and 2 cortical and medullary fibroblast subtypes (Figure 6d). Hence, these studies showed age-related transcriptomic changes across distinct NETS subtypes of *Tbx1*^*+/-*^ *Crkl*^*+/-*^ mice.

Although there was only a very small number of NCCs present at postnatal day 0 in either wild type or *Tbx1*^*+/-*^*Crkl*^*+/-*^ mice, NCCs migration and differentiation are known to be impaired in 22q.11.2 Deletion Syndrome (*46*). It is therefore possible that the alterations within the NETS compartment observed in *Tbx1*^*+/-*^ *Crkl*^*+/-*^ mice may be driven by aberrant differentiation of NCCs into mesenchymal cells, particularly perivascular cell types (*31*).

### Neural crest cells differentiate into perivascular cells in the human prenatal thymus

To extend the analysis of the thymus stroma to human tissue, we used single NETS nuclei to investigate their gene expression profiles and correlated these to chromatin accessibility. For this purpose, we employed a multiomics analysis that investigated 528 individual non-TEC thymic stroma nuclei isolated from two donors at 14- and 17-weeks post-conception (Fig. S7 and S8). This analysis identified seven distinct cell clusters, corresponding to NCCs (NCC-I and NCC-II), capsular fibroblasts, VSM, endothelial cells, medullary fibroblasts and pericytes. The frequency of cells in cluster 6 increased from 14- to 17-weeks post-conception, whereas the frequency of those in clusters 3 and 5 decreased within that time span (Fig. S9). We found the expected gradients in *PDGFRA* and *PDGFRB* expression across clusters composed of fibroblasts, pericytes and VSM (Fig. 7a) and *PECAM1* expression in endothelial cells (Fig. 7b).

**Fig. 7.**
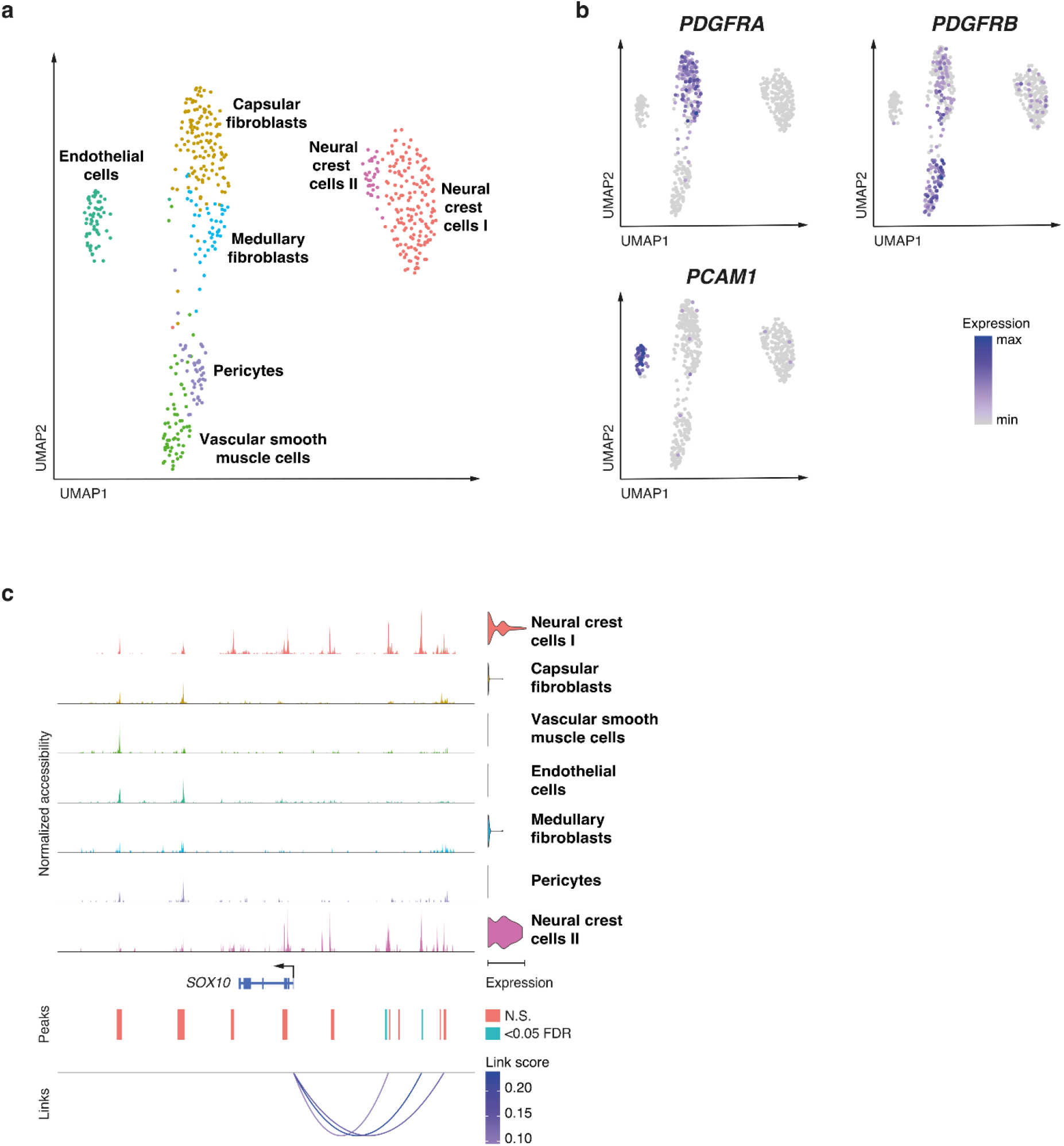
Single nuclei multiomics analysis of human thymic NES demonstrates diverse cell populations. **(a)** UMAP plot of 528 nuclei showing joint projection of transcriptomic and chromatin accessibility data. **(b)** UMAP plot showing marker gene expression for *PDGFRA, PDGFRB* and *PECAM1*. **(c)** Links plot of the *SOX10 locus* showing chromatin accessibility data, *SOX10* gene expression (violin plot on right), accessible chromatin peaks (blue peaks were significant for at least one cell type at FDR<0.05) and the correlations between chromatin accessibility and *SOX10* gene expression (shaded by the strength of each link). Significance of motif activity and chromatin accessibility were calculated using likelihood ratios, correcting for the size of chromatin accessibility libraries. P-values were adjusted for multiple hypothesis testing using Benjamini-Hochberg correction.

Given the dynamic changes in NCC frequency in the thymus over development and the changes observed in NCC-derived perivascular structures in *Tbx1*^*+/-*^*Crkl*^*+/-*^ thymi, we focused further on the chromatin and transcriptomic landscape of the NCC subclusters. NCC development is understood as a stepwise series of bifurcating cell fate decisions that lead to multiple cell identities and traits (*47*). Once specified in their fate, NCCs undergo an epithelial-to-mesenchymal transition and migrate throughout the embryo. NCC-II showed genome-wide high accessibility for sequences with SOX10 transcription factor binding motifs whereas the NCC-I displayed only an intermediate degree of accessibility for this motif (Fig. S10).

To further investigate the dynamics of SOX10 activity, we integrated gene expression profiles with chromatin accessibility in single cells to identify key enhancer-TSS interactions driving *SOX10* expression in human NCCs (Fig. 7c). We identified two sequences, which corresponded to two upstream orthologous enhancer elements (called U2 and U3) previously implicated in the control of *SOX10* expression in mouse NCCs and their progeny (*48*). Having identified that *SOX10* regulation was shared throughout the two NCC subclusters, we further examined heterogeneity amongst these NCCs to establish whether these could represent NCCs in different states of differentiation.

The comparative analysis of the gene expression profiles among the two NCC subclusters revealed an enrichment of gene pathways associated with cellular motility and vascular development for NCC-I (*e*.*g. BCL2, FGF13* and *RHOJ*; Fig. S11a). In contrast, NCC-II was enriched for gene pathways related to neuronal development and thus marked cells with gene expression profiles characteristic of bona fide NCC (*e*.*g. NES, NCAM2* and *GRID2*; Fig. S11a). This difference in gene expression profiles suggested that NCC-I may constitute a population of NCCs differentiating into mesenchymal and perivascular cell types. In support of this notion, NCC-II showed in comparison to NCC-I a significantly higher expression of *TFAP2A*, a key transcription factor in early NCC development, whereas *NR2F2*, a transcription factor involved in NCC migration, was most highly expressed in NCC-I (Fig. S11b) (*49*). Hence, the multiomics analysis of NES cells in human thymi captured the process of NCC differentiation into other cell types, a finding that could not be observed in mouse thymic stromal samples analyzed in this study.

## Discussion

Stromal cells with separate functions emerge from all germ-layers during development to populate organs where they instruct the tissue’s essential activities, for example via the differential production of extracellular matrix components, the release of growth and differentiation factors, and the creation of signalling niches that provide critical molecular cues (*50*). In addition to cross-organ communalities, stromal cells with seemingly identical phenotypes also display a heterogeneity both within and across tissues as revealed by dissimilarities in transcripts encoding pathway elements, transporters and cell-surface markers (*51*). Previous studies of the thymic stroma in both mice and humans could identify only a limited number of phenotypically distinct thymic mesenchyme subtypes (*13, 37, 52*) despite the cells’ acknowledged roles as critical components in maintaining tissue structure and TEC function (*7*).

Employing gene expression profiles at single cell resolution, we now show an unprecedented heterogeneity among thymic mesenchymal cells and identify dynamic changes in the frequency of these cells across a large range of developmental stages. Notably, the observed diversity is not replicated using flow cytometry as separate since transcriptionally defined stromal subtypes display identical phenotypic features due to a limited number of suitable cell surface markers. This limitation has hindered a comprehensive understanding of how non-epithelial thymic stromal cells contribute to local tissue microenvironments which control discrete stages of intrathymic T cell differentiation.

Early in development, the thymus stroma is mostly composed of cells belonging to the NETS and, with the notable exception of NCC, continue to structure the scaffold also in adult mice where they contribute together with TEC to the non-haematopoietic stroma. We identify within the NETS separate cell types as decoded by unique transcriptional fingerprints, including endothelial cells, vascular mural cells, NCCs, mesothelial cells and fibroblasts. Among the fibroblast population, at least 11 distinct capsular and medullary subtypes are recognized, thus largely extending the previously identified heterogeneity defined mostly by phenotypic markers and bulk RNA sequencing (*13, 37, 52*). These subtypes display dynamic changes in their relative representation over time and demonstrate RNA splicing patterns that identify CapFb1a, CapFb2b and MedFb1a as fibroblast subtypes with precursor potential and CapFb3 to originate from mesothelial cells as this fibroblast subtype continues to expresses several mesothelium-specific biomarkers, including *Wt1, Cxcl13* and *Rspo1* (Fig. S12; ref. (*14*)).

Cells with gene expression profiles typical of arterial, capillary and venous vasculature are detected already at embryonic day E12.5 when the colonization of the thymus by haematopoietic precursor cells has been initiated independent of an established vasculature (*53*). After E15.5, the frequency of pericytes and vascular smooth muscle cells increases which coincides with the histological evidence of vessel formation. The gate keeper molecules P-selectin, ICAM-1, VCAM-1 and CCL25 enable the entry of T cell precursor into the thymic microenvironment and we find these molecules expressed by all thymic endothelia in the postnatal thymus (*54*). The expression level of the adhesion molecules increases parallel to the age of the mouse but differs between distinct anatomical sites along the vasculature (Fig. S13). This expression pattern specifies that T cell precursors enter the thymic microenvironment via postcapillary venules in a gated and temporally-controlled way (*55*) and that the efficiency of this process may differ between developmental stages. The further development of these haematopoietic cells is regulated by membrane bound Kit ligand which we find expressed by all endothelia and thus also at those anatomical locations where haematopoietic precursors enter the thymus microenvironment (Fig. S12).

Endothelial cells also regulate in a non-redundant fashion the egress of mature thymocytes via the expression of sphingosine-1-phosphate (S1P) lyase (encoded by *Sgpl1*), lipid phosphate phosphatase 3 (*Pllp3*) and spinster homologue 2 (*Spns2*) that modify SIP availability and enable the molecular transport, respectively (*56-58*). Transcripts for *Ppl3* and *Spns1* are detected in all endothelial cell types even paradoxically at a time of development when T cell export has not yet commenced (Fig. S13). This expression pattern designates the anatomical site from where thymocytes can exit and indorses the molecular mechanism by which this process is controlled (*59*).

The NETS collectively promotes the proliferation and differentiation of TEC either indirectly via ligands that engage for example the platelet-derived growth factor receptor-alpha (PDGFRα) (*60*) or directly via different signalling ligands, including Wnts, BMP4, Fgf7 and Fgf10 (*3, 61, 62*). For example, BMP is expressed by CapFb1c together with Fgf10 and upregulates FOXN1, a transcription factor indispensable for TEC differentiation and function (*61-63*). Wnt4 which also stimulates the up-regulation of FOXN1 both in an auto- and paracrine way is not expressed by any of the identified thymic fibroblast subtypes but detected in mesothelial cells, thymocytes and epithelia within the thymus (*62, 64*). Moreover, three of the four capsular fibroblast subtypes express a range of Wnt ligands albeit none that had previously been implicated in stimulating FOXN1 expression. The transcriptome of individual thymic fibroblast subtypes also infers that Wnt-mediated signals are furthermore either positively modulated by R-Spondin 1 secreted by mesothelia or negatively delimited by Kremen and Dickkopf-1 which are expressed by other cellular components within the stroma (*65, 66*). Moreover, CapFb4 fibroblasts which express endosialin (*CD248*) have previously been implicated in the maintenance and regeneration of TEC (*67*).

A major feature of fibroblasts is their capacity to express extracellular matrix components which form scaffolds that differ regionally in their composition, shape, biophysical characteristics and functions (*50*). The heterogeneity and distinct spatial distribution of individual stromal cells therefore accounts for the diverse properties of the extracellular matrix with collagen expression restricted to capsular fibroblasts and transcripts for laminins detected more widely across capsular and medullary fibroblast subtypes (Fig. S3). Patterning of the extracellular matrix is critical in supporting thymic organogenesis, suggesting that thymic mesenchymal diversity will likely have a broad impact on thymic *in vivo* function (*8*).

Single cell sequencing of human thymus tissue identifies a heterogeneity among NETS that is similar to the variance observed in mice and thus constitutes a trans-species phenomenon (*37, 52*). Surprisingly, a relatively large population of NCCs is still detected at 14- and 17-weeks post-conception, i.e. at a time when thymus morphogenesis has ended and full function has been attained (*68*). This finding thus contrasts the results observed in mouse tissue at a corresponding developmental stage since the thymus of mice largely lacks NCCs as early as E16.5. This incongruity suggests that contrary to mice human NCCs have a more enduring role in shaping the NETS compartment. The analysis of human thymus tissue at late foetal stages of thymus organogenesis reveals two distinct NCC subtypes each with a seemingly different developmental potential, bespoke chromatin accessibility and proliferation dynamics. One of the two NCC subtypes, i.e. NCC-I, represents a subtype that is poised to adopt a mesenchymal and perivascular cell fate. Notably, the corresponding cell type is not identified in the mouse thymus. This suggests that the transition between neural crest-derived cells and perivascular cells in mice must either be rapid and profound, or, alternatively, occurs at an embryonic time-point that is not captured in our dataset as the lineage mapping of NCCs has previously identified their differentiation into VSM and pericytes(*12, 31*).

The murine model of 22q11DS reveals major quantitative and qualitative changes in the thymus stroma which for the first time are highlighted by a decreased proportional representation of several TEC subtypes and mesenchymal cells, the latter including a selection of capsular and medullary fibroblasts and the NCC-derived pericytes and VSM. However, *Tbx1* and *Crkl* transcripts are only solidly detected in a selection of mouse thymus fibroblast subtypes at the earliest stages of organogenesis and are notably absent in thymic NCCs. Indeed, NCCs retain their relative frequency in the presence of compound *Crkl* and *Tbx1* haploinsufficiency. Hence, the observed modifications in vascular mural cells are not the result of a reduced thymic NCC frequency or changes in *Crkl* and *Tbx*-controlled gene expression in these cells having migrated to the thymus. Rather, our data suggests that they are the consequence of altered TGFβ receptor mediated signalling in pericytes, VSM and likely their immediate precursors as this requires the involvement of Crkl, with *Tgfb1* expressed by endothelial cells and *Tgfb3* by capsular fibroblasts (Fig. S14) (*69*). Yet, the absence of normal Crkl-dependent signalling in different haploinsufficient fibroblast subtypes may in addition and indirectly impair pericytes and VSM development, thus arguing for a hitherto unexplored aspect of intra-thymic cellular crosstalk.

Overall, our findings have identified previously underappreciated levels of cellular heterogeneity and developmental dynamics within the non-TEC thymic stromal compartment. Cellular diversity within this compartment is present both in murine and human thymic development, but shows clear trans-species differences worthy of further investigation. Many of these cell populations are disrupted in 22q11.2 DS, a syndrome known to cause defective thymic organogenesis and function. Further work should focus on identifying the precise function of each fibroblast cell subpopulation, along with their contribution to overall thymic development and function.

## Materials and Methods

### Mice

All mice were maintained under specific pathogen free conditions and according to United Kingdom Home Office regulations and federal regulations and permissions, depending on where the mice were housed. Wild-type C57BL/6 mice originated were bred in-house. A mouse line carrying a germ-line Crkl null allele (Crkl^tm1d(EUCOMM)Hmgu^/ImoJ) was generated with Cre-mediated recombination in the epiblast by crossing the Crkl-flox mice (Crkl^tm1c(EUCOMM)Hmgu^/ImoJ) (*70*) with Meox2 Cre knock-in strain (*71*), followed by backcrosses with wild-type C57BL/6 mice to segregate out Meox2Cre. Tbx1^lacz/+^ (*72*) mice were obtained from Prof. Antonio Baldini via Prof. Peter Scambler at University College London. Tbx1 and Crkl compound heterozygous mice (Tbx1^+/-^Crkl^+/-^) were generated by crossing between Tbx1^+/-^ males with Crkl^+/-^ females. Embryos of specific embryonic ages were obtained through timed mating where the presence of vaginal plug was defined as embryonic day (E)0.5.

### Isolation of mouse thymic stromal cells and preparation for flow cytometry

Thymic cell suspensions were obtained via enzymatic digestion of thymic lobes using Liberase (Roche) and DNaseI (Roche). To enrich for non-haematopoietic stromal cells in thymic digests from adult mice, cell suspensions were counted and stained with anti-CD45 microbeads (Miltenyi Biotec) for 15 min on ice, before negative selection using the AutoMACS (Miltenyi Biotec) system. Enriched samples or non-enriched samples were then stained for cell surface markers for 30 min at 4°C. For intracellular staining, the Foxp3 Transcription Factor Staining Buffer Kit (eBioscience) was used according to the manufacturer’s instructions. Combinations of UEA-1 lectin (Vector Laboratories) labelled with BV605 and the following antibodies were used to stain the cells: TER-119::BV421 (BioLegend), CD45::AF700 (30-F11, BioLegend), EpCAM::PerCPCy5.5 (G8.8, BioLegend), Ly51::PE (6C3, BioLegend), CD80::PECy5 (16-10A1, BioLegend), CD26::PECy7 (H194-112, BioLegend), MHCII::APCCy7 (M5/114.15.2, BioLegend), MHCII::BV421 (M5/114.15.2, BioLegend), CD31::AF488 (MEC13.3, BioLegend), podoplanin (gp38)::AF647(PMab-1, BioLegend). DAPI or the LIVE/DEAD Fixable Aqua Dead Cell Stain Kit was used (Thermo Fisher Scientific) for the assessment of cell viability. After staining, cells were acquired and sorted using a FACS Aria III (BD Biosciences) and analysed using FlowJo v10 and GraphPad Prism 8. Statistical analyses were performed using t-tests, with correction for multiple comparisons where appropriate. A p-value or the adjusted P-value of ≤ 0.05 was considered statistically significant.

### Immunofluorescent microscopy for extracellular matrix proteins

Thymus from a 5-weeks-old female WT C57BL/6J mouse was used. The standard procedure for immunofluorescence on tissue sections was described here (https://www.biorxiv.org/content/10.1101/2021.03.21.436320v1). Briefly, organs are collected in PBS, fixed in 4% paraformaldehyde overnight at 4°C on a rotating shaker. Organs were then washed in PBS, and lobes separated for the next steps. Paraffin infiltration was done using a Tissue-Tek VIP 6 AI Vacuum Infiltration Processor (Sakura). Lobes were then embedded in paraffin and 4μm sections cut with a Hyrax M25 microtome (Zeiss).

Before immunostaining, de-waxing and antigen retrieval in citrate buffer at pH6.0 (using a heat-induced epitope retrieval PT module, ThermoFischer Scientific) were performed. Sections were then blocked and permeabilized for 30min in 1% BSA, 0.2% Triton X-100 in PBS and blocked for 30min in 10% donkey serum (Gibco) in PBS at RT. Sections were incubated with primary antibodies overnight at 4°C in 1.5% donkey serum in PBS. Sections were washed twice in 1% BSA, 0.2% Triton X-100 in PBS and incubated with secondary antibodies at RT for 45min. Finally, sections were washed twice in 0.2% Triton X-100 in PBS and mounted with Fluoromount-G (SouthernBiotech). Pictures were acquired with a CCD DFC 3000 black and white camera on an upright Leica DM5500 scanning microscope.

Antibodies: goat anti-Fibronectin (Santa Cruz, sc-6953, 1/250); rabbit anti-Collagen 1 (Abcam, ab21286, 1/250); Donkey anti-goat Alexa 488 (ThermoFischer Scientific, A-11055, 1/500); Donkey anti-rabbit Alexa 647 (ThermoFischer Scientific, A-31573, 1/500). Nuclei staining: DAPI (Sigma-Aldrich, 1μg per ml).

### Immunofluorescent microscopy for CD26 and podoplanin

Freshly isolated thymic lobes were frozen in OCT compound (Tissue-Tek) and cryosectioned at a thickness of 10 μm. Tissue sections were fixed with ice cold acetone for 5 min and blocked with Avidin/Biotin Blocking Kit (Vector laboratories) and Protein block (Protein block (Dako) according to manufacturer’s protocol. Tissue sections were then incubated with primary antibodies at 4 °C overnight: rabbit anti-mouse CD26 (DPP4) (EPR5883(2), Abcam) and biotin anti-mouse podoplanin (8.1.1, Biolegend). Secondary antibody staining was performed at room temperature for 30 min with anti-rabbit:AF488 (Invitrogen) and streptavidin-AF555 (Invitrogen)). Nuclei were stained with Hoechst 34580 in PBS (according to manufacturer’s protocol). Sections were mounted with ProLong Gold Antifade Mountant (Thermo Fisher Scientific) and acquired using an LSM700 confocal microscope (Carl Zeiss AG). Image analysis was performed with ImageJ software (Rasband WS, ImageJ, US National Institutes of Health, Bethesda, Md).

### Single cell RNA sequencing

Total thymic non-epithelial stromal cells (Live Ter119-CD45-EpCAM-) thymic cells from E12.5, E13.5, E16.5, P0 and 4-week-old wild type mice were sorted and kept on ice before they were counted. 18,000 cells per sample were loaded onto a Chromium Single Cell B Chip (10x Genomics) followed by library preparation using Chromium Single Cell 3’ solution (10x Genomics) and sequencing by NovaSeq6000. (28+98) (Illumina). For the Tbx1^LacZ/+^Crkl^+/-^ dataset, total non-haematopoietic stromal cells (Live Ter119-CD45-) from P0 Tbx1^LacZ/+^Crkl^+/-^ (n=3) and their wildtype littermates (n=3) were sorted and fixed using RNAprotect Cell Reagent (Qiagen) for storage before sample submission to the Oxford Genomics Centre, where all downstream steps were performed including 10x Genomics Chip loading, library preparation and sequencing.

### Single cell RNA sequencing analysis

Sequencing reads were processed using Cell Ranger (version 3.1.0). Cells were retained for downstream analysis if there was expression of >1,000 genes, <5% of UMIs mapped to mitochondrial genes, cells were called as singlets by DoubletFinder, and cells did not cluster into *Ptprc* (CD45)-expressing clusters or other contaminant clusters (such as thymic epithelial cells or clusters present only in one replicate) (*73*). Seurat was used to remove batch effect between samples using canonical correlation analysis-based integration (*74*). Cells were projected into two-dimensional space using Uniform Manifold Approximation and Projection (UMAP). Clusters were called using a resolution of 0.8 and cell label transfer between datasets was undertaken using Seurat. Differential analysis between clusters used Wilcoxon-rank sum testing and over different ages used the Kruskal-Wallis analysis of variance. P-values were corrected for multiple hypothesis testing using the Benjamini-Hochberg method. GENIE3 and RcisTarget were used to identify gene-regulatory networks on highly variable genes expressed in at least 5% of cells with subsequent module expression calculated using Seurat (*75*). RNA velocity analysis was undertaken using Velocyto and scVelo (*76, 77*). Ligand-receptor-target networks were inferred using Nichenetr, with differential expression assessed between early (E12.5/13.5) and late (E16.5) embryogenesis (*38*). Gene ontology analysis was undertaken using clusterProfiler (version 4.0.0) (*78*).

### Single nuclei multiomics of human fetal thymic stroma

Human fetal thymi, obtained from terminations of pregnancy at 14 and 17 post conception weeks were enzymatically dissociated using Liberase (Roche) and DNaseI (Roche). The resultant cell suspension was stained with the following antibodies directed against cell surface antigens for 30 minutes at 4°C: CD45::BV421 (H130, BioLegend) and HLA-DR::PE-Cy7 (L243, BioLegend); 7AAD (BioLegend) was used as a viability marker. Live CD45-MHCII intermediate-high cells were sorted in 250, 000 cell aliquots using a FACS Aria III (BD Biosciences). Samples were then processed using the 10x Genomics Multiomics ATAC (Assay for Transposase-Accessible Chromatin using sequencing) + Gene Expression kit according to the manufacturer’s protocol with some adaptations. Specifically, nuclei were isolated using a 0.1x diluted nuclei extraction buffer for 6 minutes before being captured into droplets on the 10x Genomics Chromium platform and sequenced on an Illumina NovaSeq machine.

This study of human thymic tissue has been granted ethical approval and is publicly listed (IRAS ID 156910, CPMS ID 19587).

### Multiomics analysis

Sequencing data were processed using Cell Ranger ARC (version 1.0.1). Counts and ATAC data were analysed using Seurat (version 4.0.3) and Signac (version 1.2.1) (*74, 79*). Barcodes were filtered to high quality cells (ATAC library size 1,000-100,000, RNA library size 1,000-31,622, ATAC peaks 1,000-31,622, RNA features 1,000-10,000 and proportion of mitochondrial RNA reads ≤0.15). ATAC peaks were recalled across each sample for all cells. Clusters were called on integrated RNA data used a clustering threshold of 0.8 and projected onto a joint UMAP plot of RNA and ATAC components generated using Seurat and Signac. Differential gene expression between clusters was estimated using the default method in Seurat. Differentially accessible peaks were identified using the likelihood ratio method with correction for ATAC library size. Motif activity was estimated using chromVAR with the JASPAR2020 motif dataset (*80*). RNA-ATAC links were analysed using Signac and Seurat in 50kb windows around genes of interest.

## Funding

Swiss National Science Foundation grant IZLJZ3_171050; 310030_184672 (GAH) Medical Research Council grant MR/S036407/1 (GAH, AEH)

Wellcome Trust grant 105045/Z/14/Z (GAH)

The work was generously supported by a donor of Stanford’s Center for Definitive and Curative Medicine 22q11 Deletion Syndrome Consortium (GAH)

National Institute for Health Research Oxford Biomedical Research Centre grant (GAH)

NIHR Clinical Lectureship (AEH, FD)

Health Research Bridging Salary Scheme (AEH) Wellcome Trust grant (FD)

International postdoc fellowship from the Swedish research council (Vetenskapsrådet) (SC)

The human embryonic and fetal material was provided by the Joint MRC/Wellcome Trust (grant # MR/006237/1) Human Developmental Biology Resource (www.hdbr.org).

## Author contributions

Conceptualization: AEH, SC, ML, KW, GAH

Investigation: SC, FD, SM, TH, MED

Analysis: AEH, SC

Visualization: AEH, SC, FD, IR, TH, GAH

Supervision: ML, GAH

Writing—original draft: AEH, SC, GAH

Writing—review & editing: AEH, SC, FD, SM, TH, IR, MED, OE, ML, KW, GAH

