## Supplementary figures for "Developmental dynamics of the neural crest-mesenchymal axis in creating the thymic microenvironment" (from another version of this preprint: version 1, 2021-11-08)

**This PDF file includes:**

Figs. S1 to S14

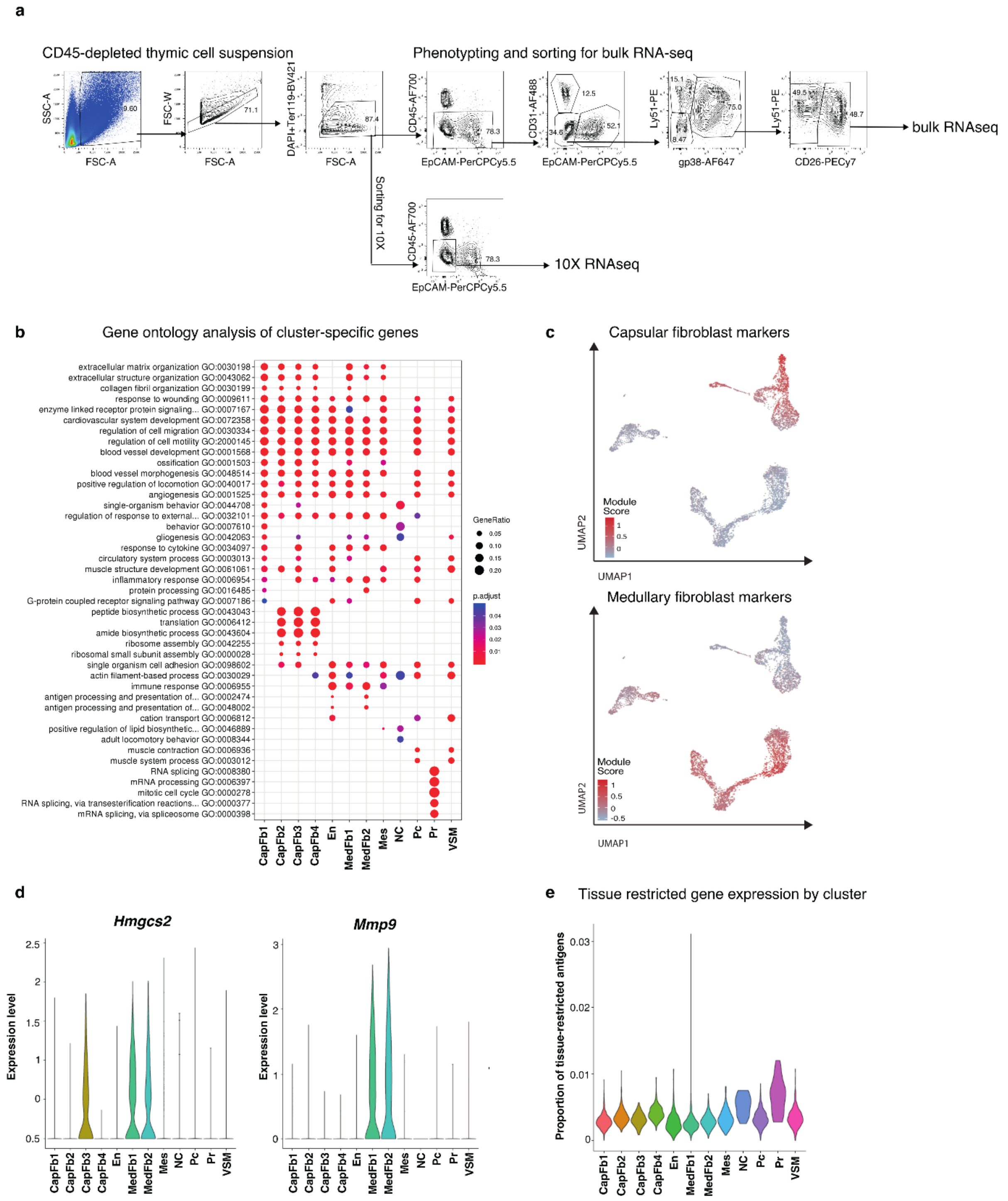


**Fig. S1.**

**Single cell phenotyping of the thymic stroma at 4 weeks old**. (**a**) Gating and sorting strategy of thymic stroma populations in 4-week-old wildtype mice **(b)** Gene ontology analysis of cluster-specific genes. (**c**) A UMAP plot of the expression of genes specific to (**i**) capsular or (**ii**) medullary fibroblasts [13]. (**d**) Violin plots of the expression of *Hmgcs2* and *Mmp9*, tissue-restricted genes specifically expressed by medullary fibroblasts. (**e**) A violin plot of the proportion of UMIs mapping to tissue-restricted antigens (defined as $\tau \geq 0.8$ in the mouse ENCODE RNA-seq dataset).

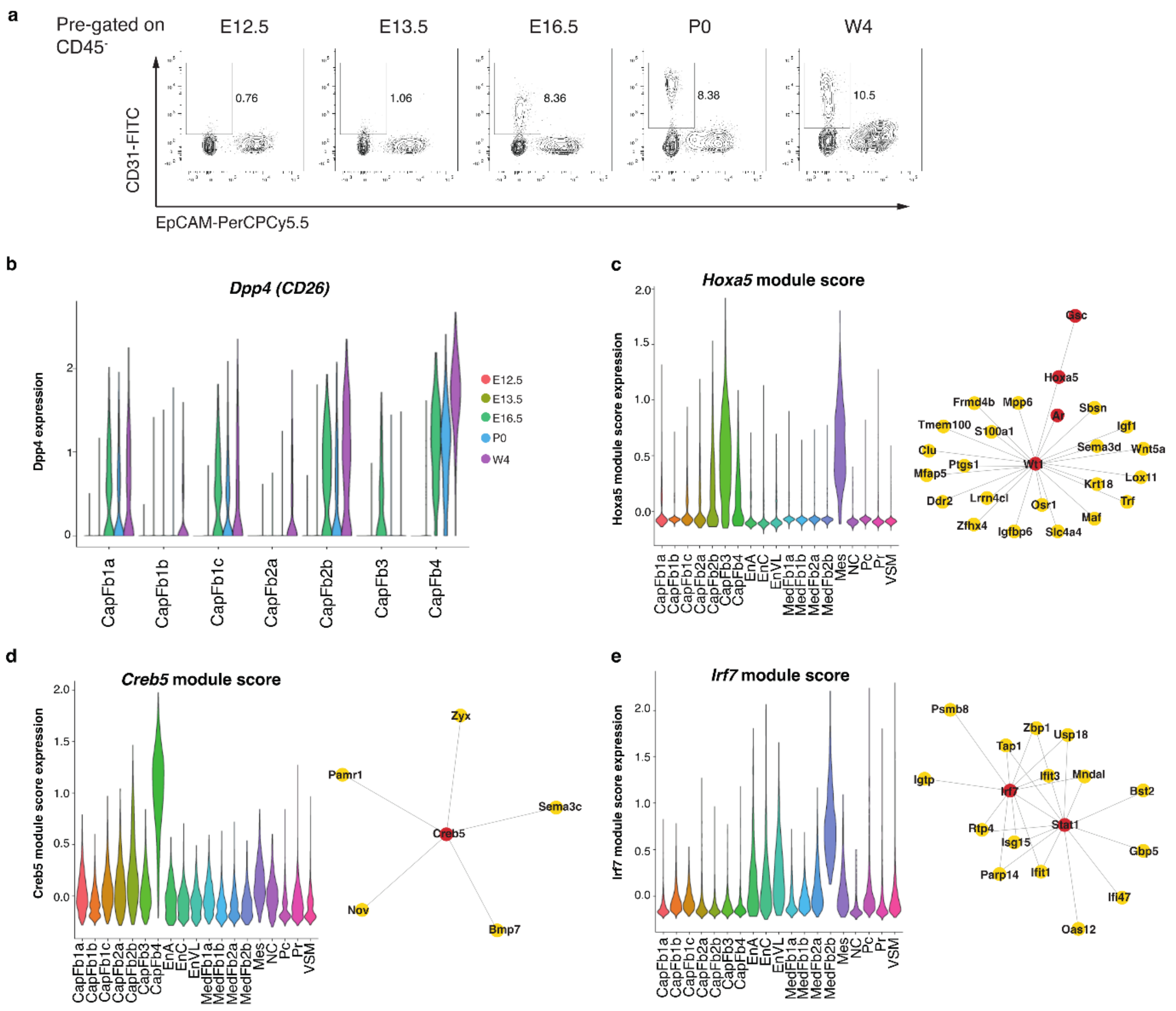


**Fig. S2.**

***Dpp4* (CD26) expression and gene regulatory network activity within the thymic stroma.**
**(a)** Representative FACS plots showing CD31+ population in E12.5, E13.5, E16.5, P0 and week 4. **(b)** Violin plot showing CD26 (*Dpp4*) expression within capsular fibroblast subsets by age. (**c**-**e**) Violin plots of gene-regulatory expression and network plots for (**c**) *Hoxa5*, (**d**) *Creb5* and **(e)** *Irf7*.

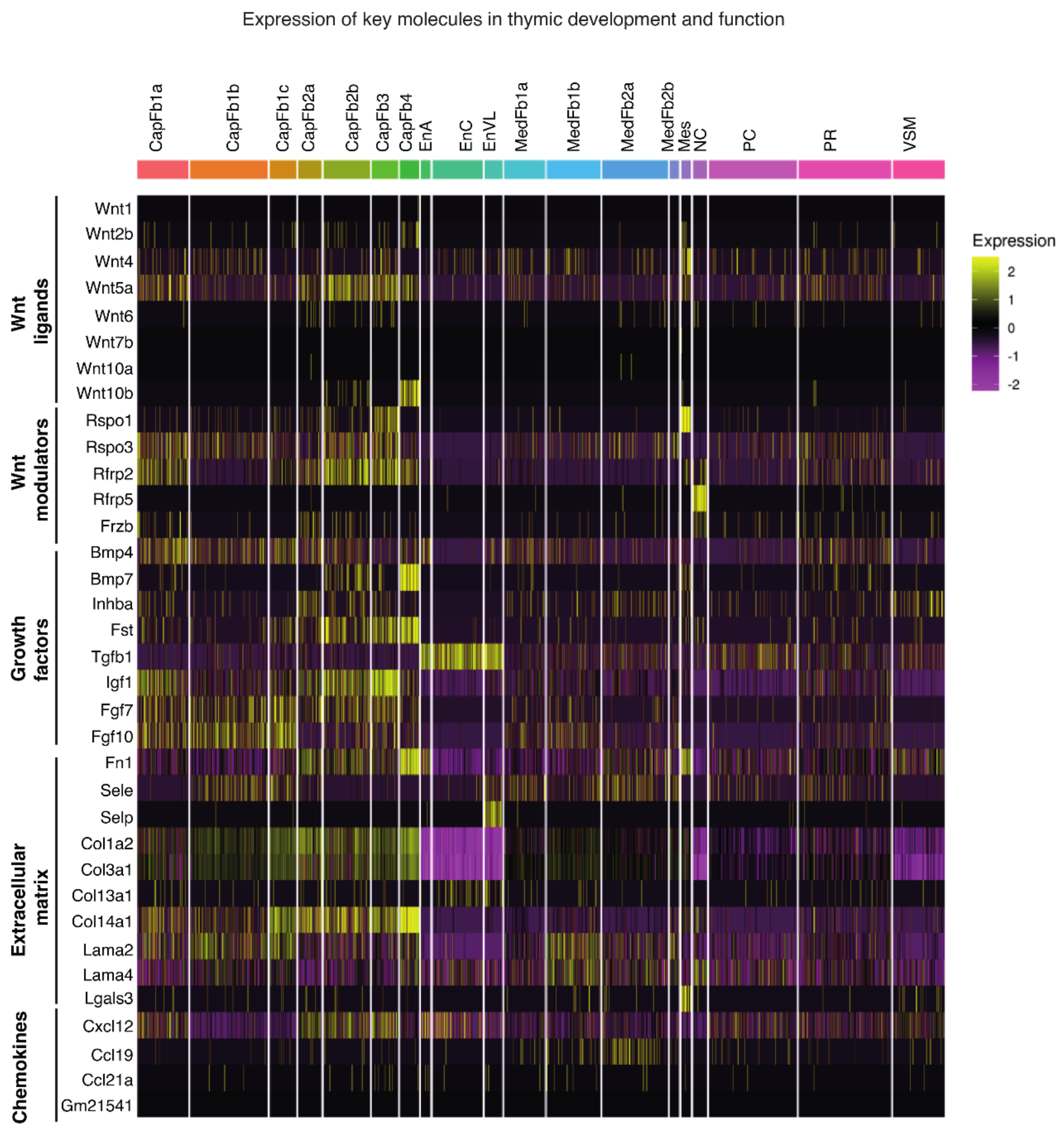


**Fig. S3.**

**Expression of key molecules in thymic development and function.** A heatmap of the scaled expression of curated transcripts across different cell subtypes. The dataset was down-sampled to 30,000 cells for visualization.

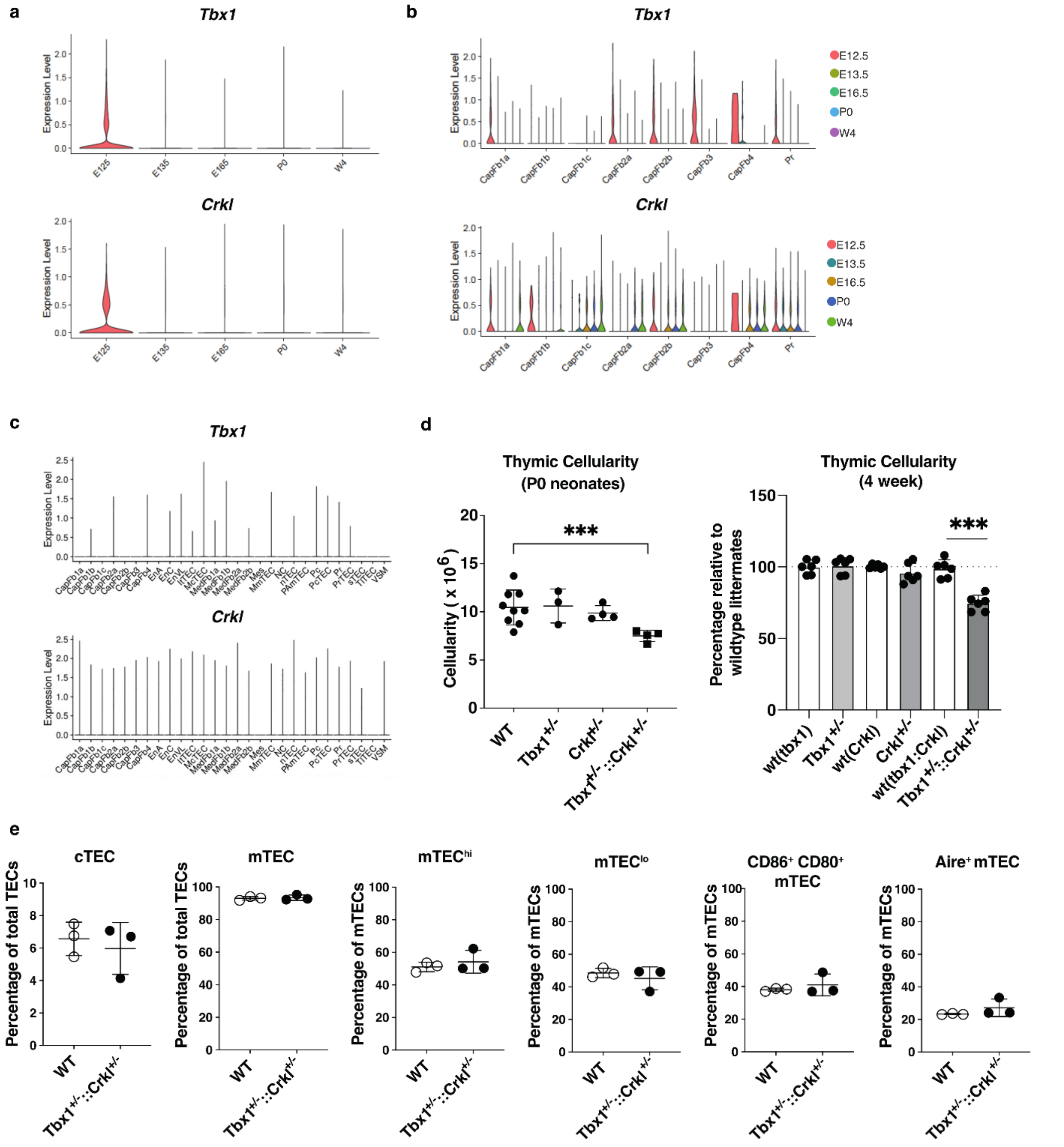


**Fig. S4.**

**Total cellularity in single heterozygous and TEC phenotypes in *Tbx1*$^{+/-}$*Crkl*$^{+/-}$ mice. (a)** Violin plot of *Tbx1* and *Crkl* expression across non-TEC thymic stroma by age. **(b)** Violin plot of *Tbx1* and *Crkl* expression split by age in those cell populations with substantial expression of *Tbx1* and *Crkl*: CapFb subsets and proliferating fibroblasts. **(c)** Violin plot of *Tbx1* and *Crkl* expression across all thymic stroma subsets including epithelial cells at P0 **(d)** Total thymic cellularity of P0 and 4-week-old *Tbx1*$^{+/-}$, *Crkl*$^{+/-}$, *Tbx1*$^{+/-}$*Crkl*$^{+/-}$ and their wildtype littermate controls. **(e)** TEC phenotype of 4-week-old *Tbx1*$^{+/-}$*Crkl*$^{+/-}$ and their wildtype littermate controls. Data shown in **(e)** from a representative experiment from at least two independent experiments.

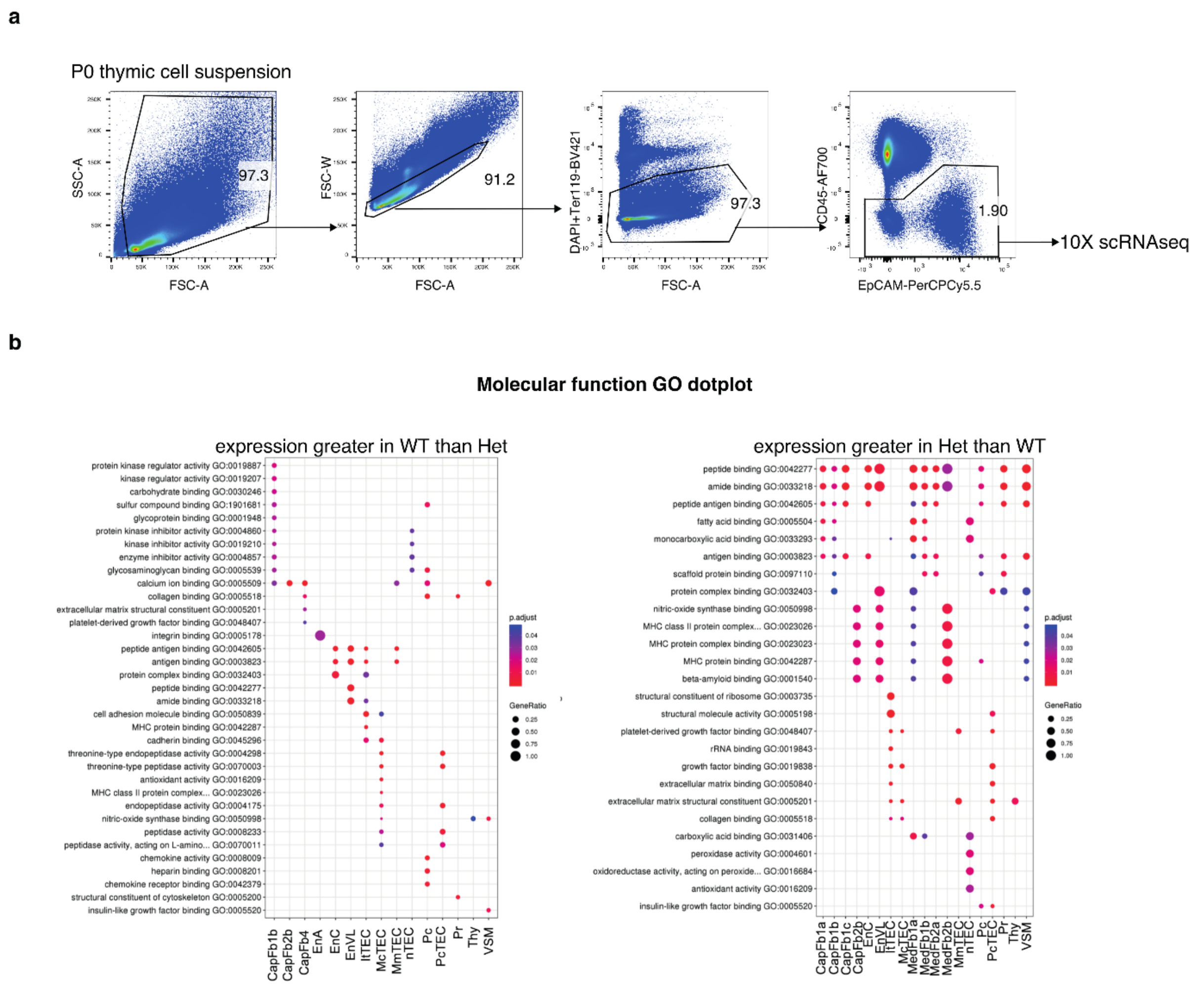


**Fig. S5.**

**Single cell RNAseq analysis of the thymic stroma of Tbx1$^{+/-}$Crkl$^{+/-}$ and wild-type mice**. (a) Sorting strategy of Ter119$^{-}$ CD45$^{-}$ total thymic non-haematopoitic stroma from P0 mice. (b) Pathway enrichment analysis (cellular components) of differentially expressed gene between *Tbx1$^{+/-}$Crkl$^{+/-}$* and wildtype thymic stromal cell subsets.

a

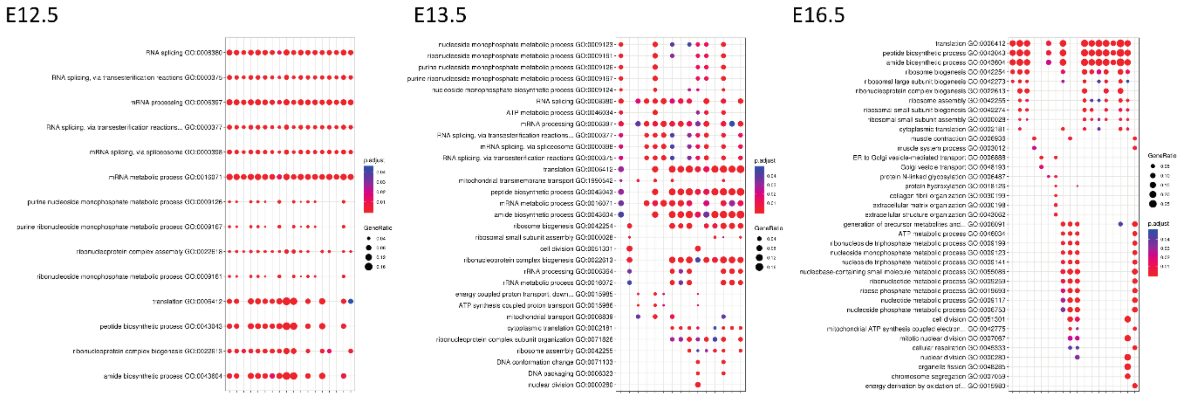


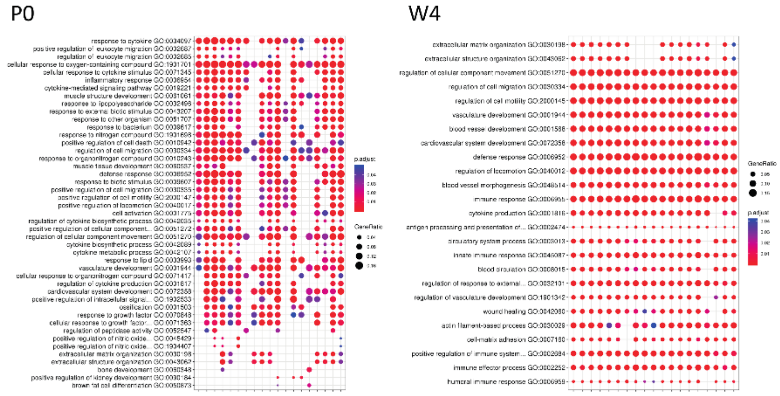


| | E12.5 | E13.5 | E16.5 | P0 | W4 |
|---|---|---|---|---|---|
| CapFb1a | X | | X | X | X |
| CapFb1b | X | | X | X | X |
| CapFb1c | X | X | X | X | X |
| CapFb2a | X | X | X | X | X |
| CapFb2b | X | | X | X | X |
| CapFb3 | X | | X | X | X |
| CapFb4 | X | X | X | X | X |
| EnA | X | X | X | X | X |
| EnC | X | X | X | X | X |
| EnVL | X | X | X | X | X |
| MedFb1a | X | X | X | X | X |
| MedFb1b | X | X | X | X | X |
| MedFb2a | X | X | X | X | X |
| MedFb2b | | X | X | X | X |
| Mes | X | X | | X | X |
| NC | X | X | X | X | X |
| Pc | X | X | X | X | X |
| Pr | X | X | X | X | X |
| VSM | X | X | X | X | X |

b

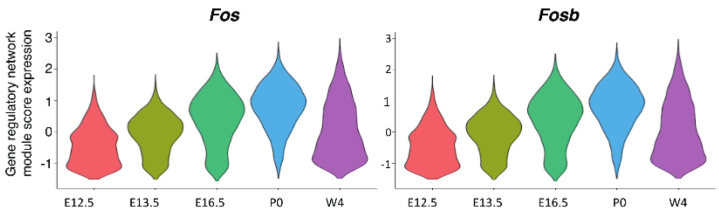


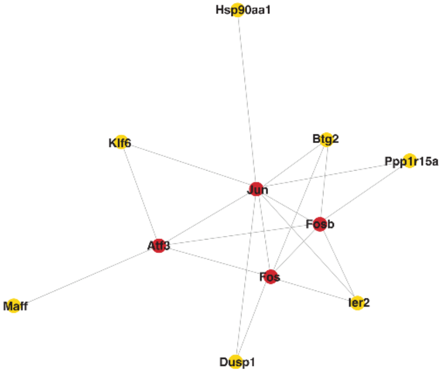


c

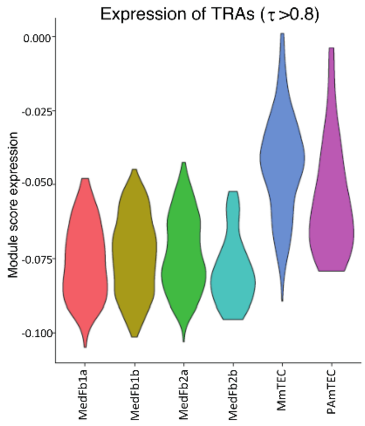

**Fig. S6.**

**Age-specific expression patterns in thymic stroma.** (**a**) GO plots of age-specific expression within each cell type. The inset table shows cell types illustrated in each plot (top to bottom in the table = left to right in the dot plots). (**b**) Violin plots of *Fos* and *Fosb* gene-regulatory network expression (top) and network plot (bottom). **(c)** Violin plot of tissue restricted gene expression within wild-type medullary fibroblasts or mTEC cells at P0.

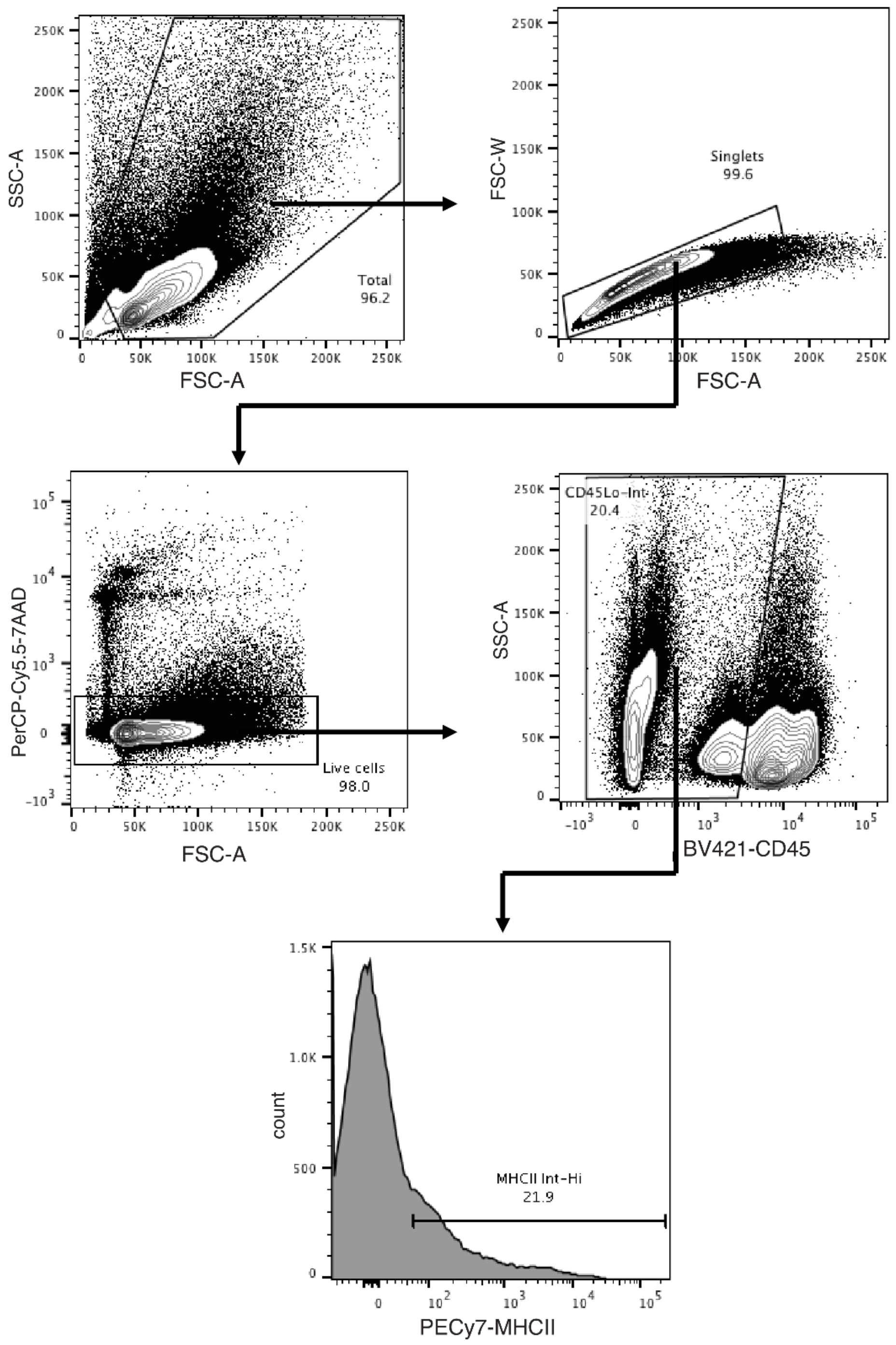


**Fig. S7.**

**FACS strategy to isolate human thymic stromal cells for multiomics analysis.**

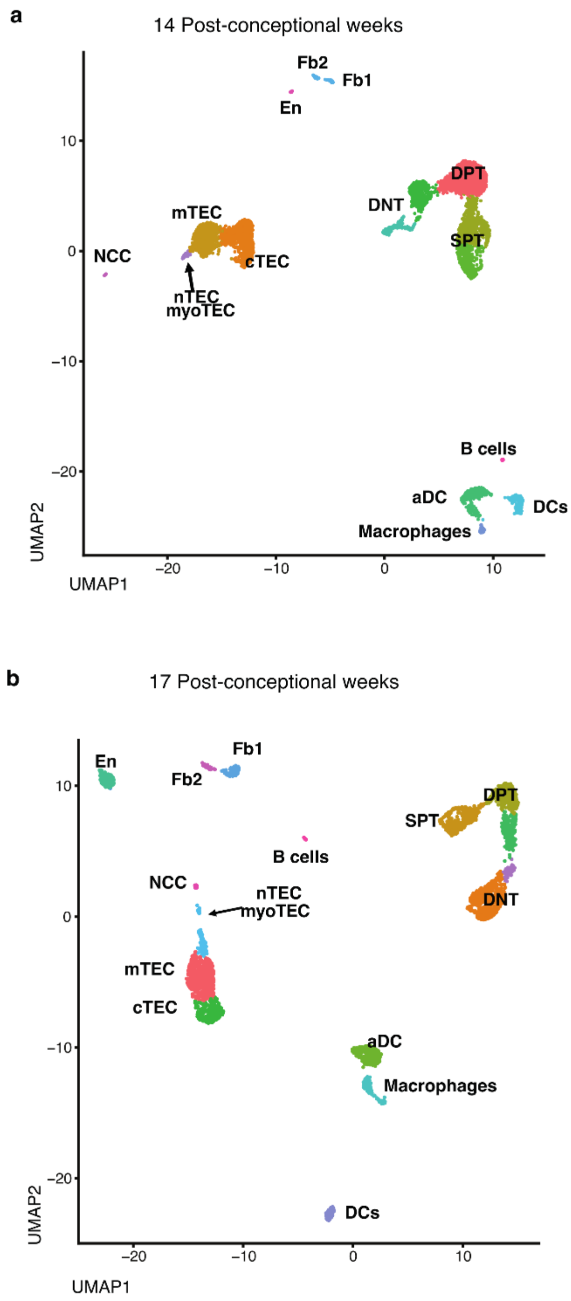


**Fig. S8.**

**UMAP projections of human thymic stroma.** UMAP plots of human thymic stromal nuclei sorted as live CD45- MHC intermediate-high at **(a)** 14 post-conceptional weeks and **(b)** 17 post-conceptional weeks. Clusters were called using the RNA and ATAC components. Cell type annotations were added manually using marker genes plus similarity to reference datasets and the orthologous datasets described earlier [13, 52]. Nuclei from the NES clusters were taken forward for further analysis. aDC = activated dendritic cells; B = B cells; cTEC = cortical TEC; DC = dendritic cells; DNT = double negative thymocytes; DPT = double positive thymocytes; En = endothelium; Fb1 = fibroblast 1; Fb2 = fibroblast 2; Mf = macrophages; mTEC = medullary

TEC; myoTEC = myoid TEC; NCC = neural crest cells; nTEC = neural TEC; SPT = single positive thymocytes.

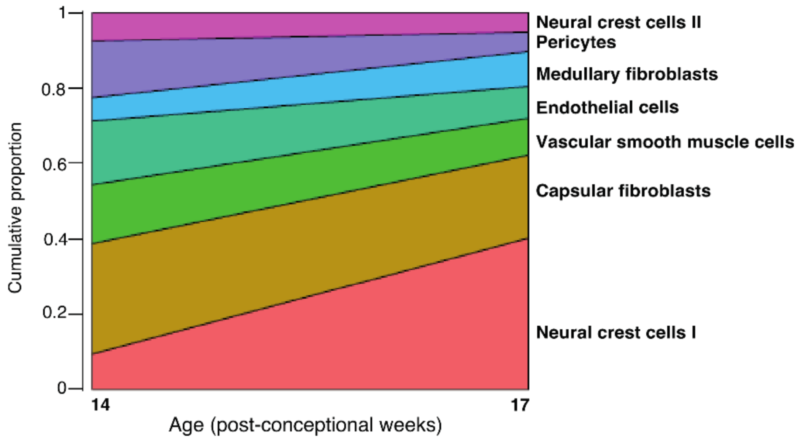


**Fig. S9.**

**Changes in human thymic NES composition between 14- and 17-post-conceptional weeks.** Cell type colours are as per Figure 7a.

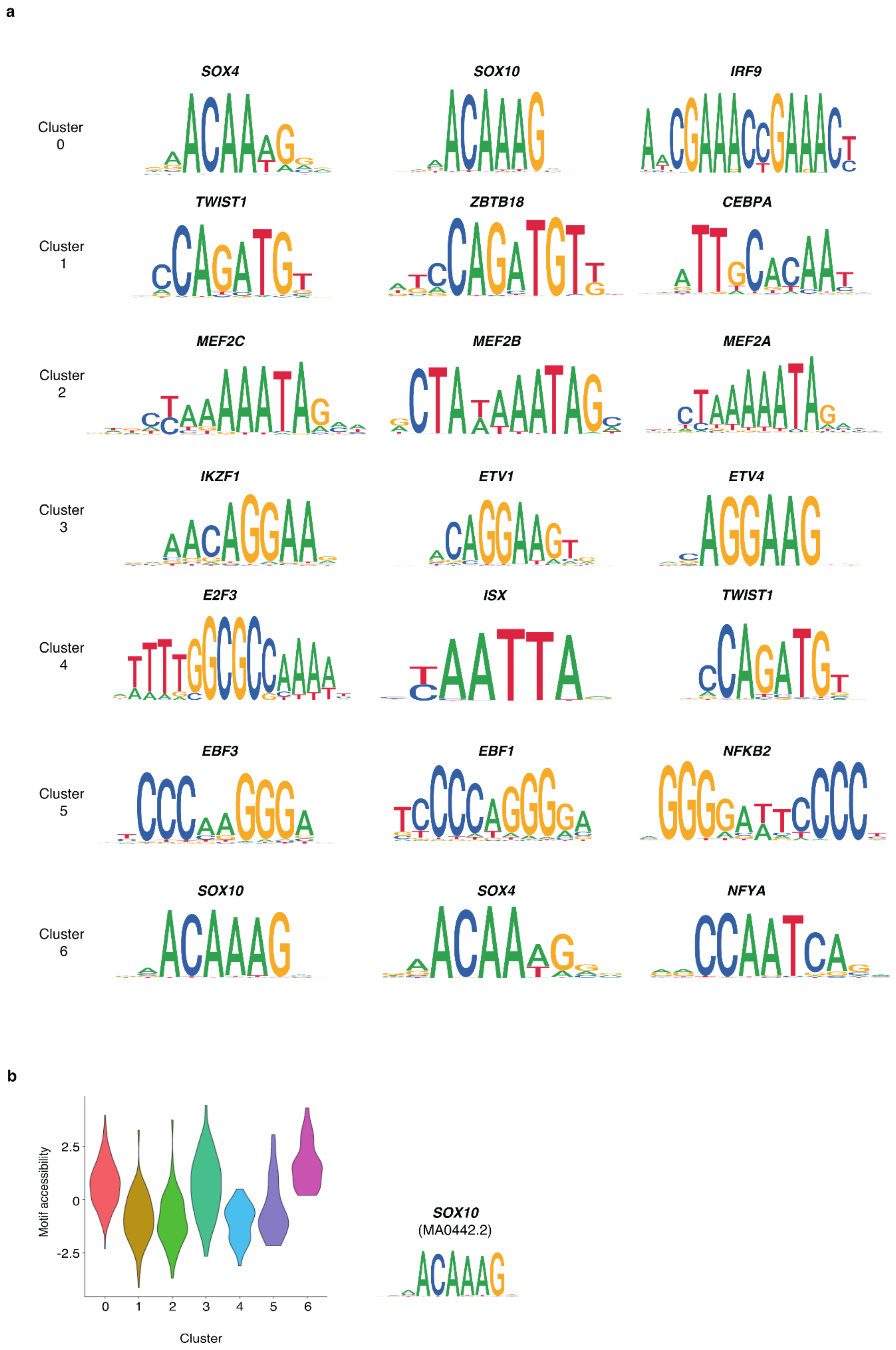
a
SOX4
SOX10
IRF9
Cluster 0
TWIST1
ZBTB18
CEBPA
Cluster 1
MEF2C
MEF2B
MEF2A
Cluster 2
IKZF1
ETV1
ETV4
Cluster 3
E2F3
ISX
TWIST1
Cluster 4
EBF3
EBF1
NFKB2
Cluster 5
SOX10
SOX4
NFYA
Cluster 6
b
Motif accessibility
2.5
0
-2.5
0
1
2
3
4
5
6
Cluster
SOX10
(MA0442.2)

**Fig. S10.**

**Motif activity across human thymic NES. (a)** Top 3 motif activity for each cluster as ranked by log fold change. **(b)** SOX10 motif activity across all clusters.

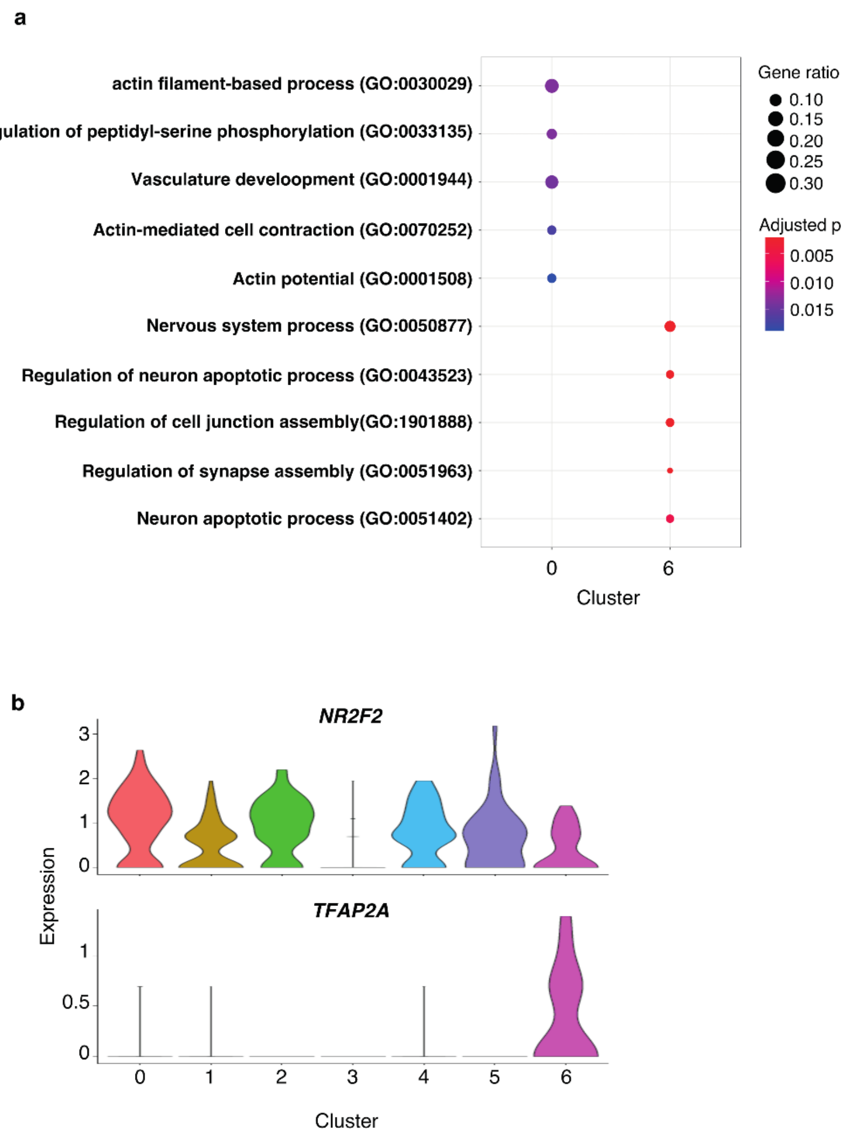


**Fig. S11.**

**A transcriptomic comparison of clusters 0 (NCC-I) and 6 (NCC-II).** (**a**) A dotplot showing significantly upregulated biological process pathways for clusters 0 (NCC-I) and 6 (NCC-II) respectively. Geneset enrichment was calculated using hypergeometic testing in clusterProfiler with adjustment for multiple hypothesis testing using Benjamini-Hochberg correction. (**b**) Violin plots of key transcription factors involved in NCC development and migration.

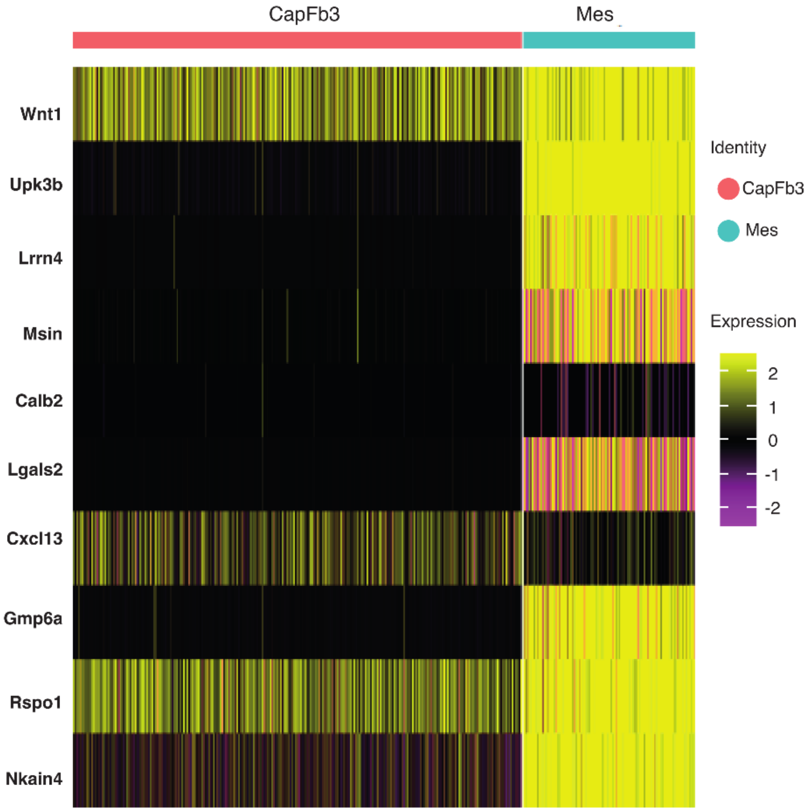


**Fig. S12.**

**Heatmap showing the expression of mesothelial cell markers within the CapFb3 and mesothelial clusters.**

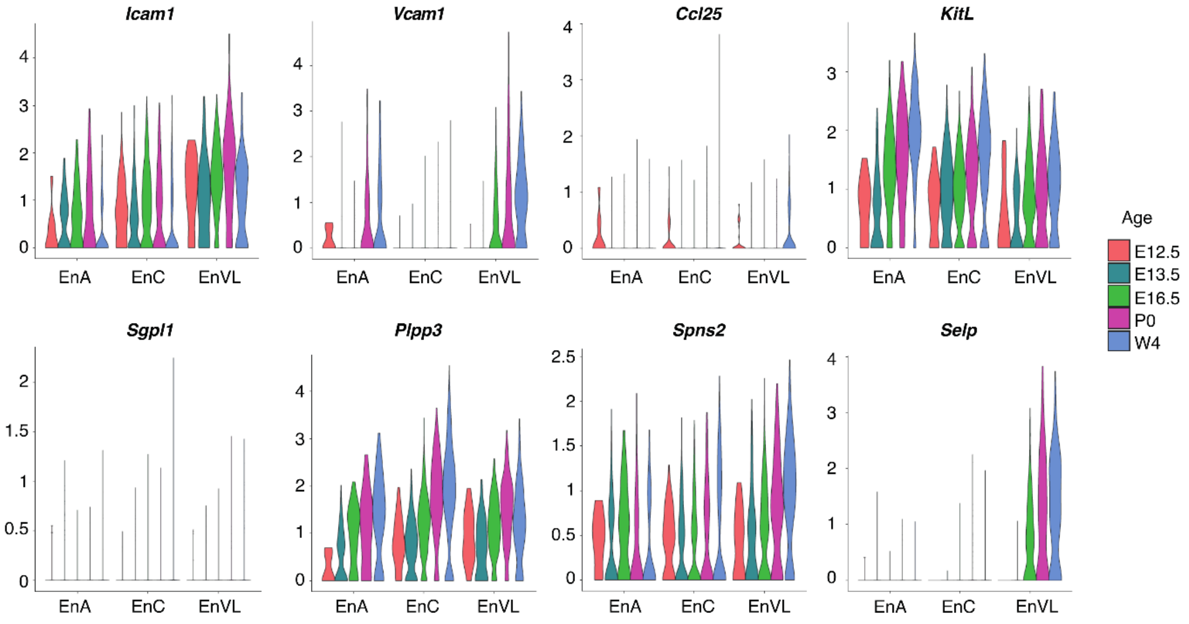


**Fig. S13.**

**Violin plots showing the expression of genes associated with endothelial adhesion or thymocyte migration within endothelial cells over different ages.**

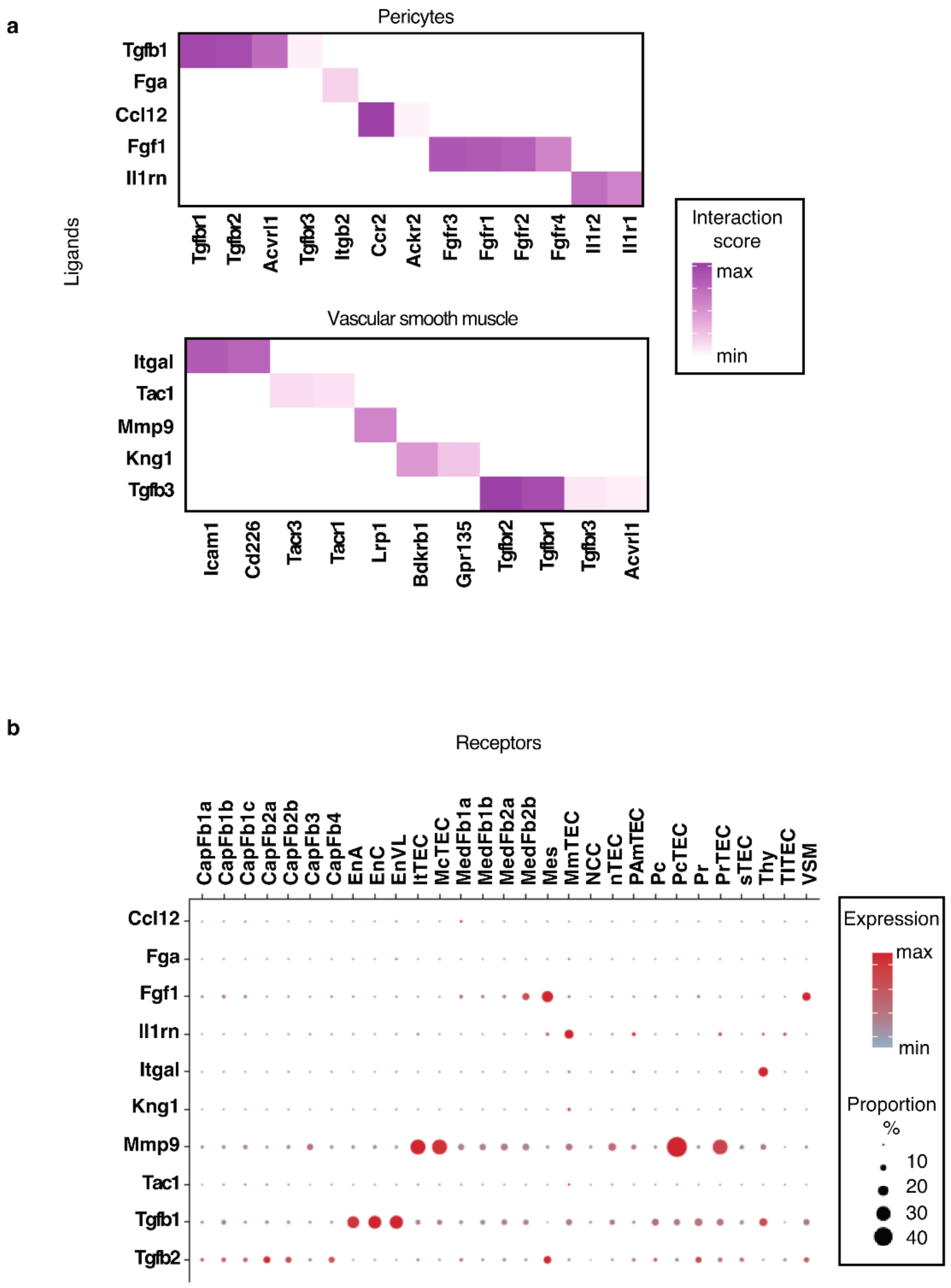


**Fig. S14.**

**Ligand-receptor signaling within perivascular cell populations in *Tbx1*$^{+/-}$ *Crkl*$^{+/-}$ mice.** (**a**) Heatmaps showing the interaction strength of the top 5 *bona fide* ligand-receptor signaling pathways driving the transcriptomic difference between *Tbx1*$^{+/-}$ *Crkl*$^{+/-}$ and wild type pericytes and vascular smooth muscle cells. (**b**) A dot plot showing the expression of these ligands within different cell populations.
